# Chronic metabolic stress drives developmental programs and loss of tissue functions in non-transformed liver that mirror tumor states and stratify survival

**DOI:** 10.1101/2023.11.30.569407

**Authors:** Constantine N. Tzouanas, Marc S. Sherman, Jessica E.S. Shay, Adam J. Rubin, Benjamin E. Mead, Tyler T. Dao, Titus Butzlaff, Miyeko D. Mana, Kellie E. Kolb, Chad Walesky, Brian J. Pepe-Mooney, Colton J. Smith, Sanjay M. Prakadan, Michelle L. Ramseier, Evelyn Y. Tong, Julia Joung, Fangtao Chi, Thomas McMahon-Skates, Carolyn L. Winston, Woo-Jeong Jeong, Katherine J. Aney, Ethan Chen, Sahar Nissim, Feng Zhang, Vikram Deshpande, Georg M. Lauer, Ömer H. Yilmaz, Wolfram Goessling, Alex K. Shalek

**Author notes:** Correspondence (A.K.S), (W.G.), (Ö.H.Y.). These authors contributed equally (Constantine N. Tzouanas, Marc S. Sherman, Jessica E.S. Shay). These senior authors contributed equally (Ömer H. Yilmaz, Wolfram Goessling, Alex K. Shalek).

## Abstract

Under chronic stress, cells must balance competing demands between cellular survival and tissue function. In metabolic dysfunction-associated steatotic liver disease (MASLD, formerly NAFLD/NASH), hepatocytes cooperate with structural and immune cells to perform crucial metabolic, synthetic, and detoxification functions despite nutrient imbalances. While prior work has emphasized stress-induced drivers of cell death, the dynamic adaptations of surviving cells and their functional repercussions remain unclear. Namely, we do not know which pathways and programs define cellular responses, what regulatory factors mediate (mal)adaptations, and how this aberrant activity connects to tissue-scale dysfunction and long-term disease outcomes. Here, by applying longitudinal single-cell multi-omics to a mouse model of chronic metabolic stress and extending to human cohorts, we show that stress drives survival-linked tradeoffs and metabolic rewiring, manifesting as shifts towards development-associated states in non-transformed hepatocytes with accompanying decreases in their professional functionality. Diet-induced adaptations occur significantly prior to tumorigenesis but parallel tumorigenesis-induced phenotypes and predict worsened human cancer survival. Through the development of a multi-omic computational gene regulatory inference framework and human *in vitro* and mouse *in vivo* genetic perturbations, we validate transcriptional (RELB, SOX4) and metabolic (HMGCS2) mediators that co-regulate and couple the balance between developmental state and hepatocyte functional identity programming. Our work defines cellular features of liver adaptation to chronic stress as well as their links to long-term disease outcomes and cancer hallmarks, unifying diverse axes of cellular dysfunction around core causal mechanisms.

## Introduction

Cells must balance their immediate viability with supporting tissue homeostasis and contributing to organismal health^1^. The hepatocytes of the liver, for example, perform wide-ranging professional functions, including nutrient metabolism, protein secretion, and chemical detoxification^2,3^. Moreover, they possess substantial regenerative capacity, helping to restore normal mass and function after acute challenges as dramatic as surgical removal of two-thirds of the liver’s mass^4^. However, this intrinsic regenerative ability can prove insufficient during chronic stress, resulting in progressive tissue damage^5,6^. For instance, chronic exposure to high caloric diets can precipitate metabolic dysfunction-associated steatotic liver disease (MASLD, formerly NAFLD/NASH; affecting ∼25% of people around the world), which in turn drives progressive fibrosis, cirrhosis, liver failure, and hepatocellular carcinoma (HCC), the second-leading cause of years of life lost to cancer^7–13^.

Epidemiological studies indicate that each successive MASLD stage associates with a progressive increase in HCC incidence^14^. However, mutations mainly accumulate after cirrhosis, but not before^15,16^: among pre-cancer liver disease patients, only patients with cirrhotic livers (but not earlier fibrosis stages) exhibited significant increases in mutation rate compared to patients with non-fibrotic livers^17^. Among a MASLD-predominant cohort, convergent somatic mutations were enriched for metabolic enzymes, but largely did not overlap with HCC driver mutations^18^. These epidemiological and mutation cohort studies raise the hypothesis that, in addition to experiencing mutational accumulation, hepatocytes may respond to chronic stress by developing progressively dysfunctional cell states that, while not solely genetically defined, are poised for tumorigenesis. Work in other organs has described environmental stressors driving non-mutational priming for longer-term dysfunction and tumorigenesis, manifesting as transcriptional and epigenetic adaptations in response to inflammation in the pancreatic and skin epithelia and high-fat diets in intestinal stem cells^19–24^.

However, prior work in MASLD has largely focused on tissue-level histology or organ-level function in the context of specific gene knockouts or immune subset depletions, or examined broad contributors to cell death, such as reactive oxygen species, unfolded protein response, or lipotoxicity^7–9,25–28^. Comparatively little is known about phenotypic changes in surviving cells and their dynamics. Outstanding questions include: which pathways and functional tradeoffs are induced with progressive exposure to environmental stressors? How do early adaptations connect to long-term consequences like tumor outcomes? And, which decision-making circuits causally mediate cellular (mal)adaptations? Knowledge of the temporal hierarchy of stress adaptations (and their accompanying disease repercussions) would help elucidate how the liver coordinates homeostatic functions while buffering stresses affecting constituent cells^29,30^. Furthermore, the discovery of cell-extrinsic and cell-intrinsic drivers of these processes could lead to novel therapeutic targets and improved patient stratification for individuals with MASLD or HCC.

Here, we examine how chronic metabolic stress drives functional tradeoffs between cellular identity and homeostatic function among hepatocytes to precipitate cancer-associated states. We conduct longitudinal single-cell multi-omics analyses of a diet-only mouse model of chronic stress via metabolic overload, spanning the stages of early steatosis to spontaneous tumorigenesis. With these resources and extensions to human MASLD/HCC cohorts, we define the progression of hepatocyte adaptation, including upregulation of early developmental markers, anti-apoptotic/pro-survival effectors, and WNT signaling. These shifts occur at the expense of core identity and professional functions, resulting in reduced expression of lineage-determining transcription factors, rate-limiting enzymes, and immunomodulatory secreted proteins. Through the development of a computational framework to discover putative regulators of disease-associated gene programs and experimental genetic perturbations (human *in vitro* and mouse *in vivo*), we validate RELB, SOX4, and HMGCS2 as causally driving hepatocyte dysregulation and inducing early shifts towards development- and cancer-associated states. Our results define principles of cellular response to chronic stress in non-transformed liver tissue and connect them to cancer-associated sequelae, suggesting avenues by which initial stress adaptations perturb cellular states, priming them for long-term tissue dysfunction and disease outcomes.

## Results

### Long-term high fat diet in wild-type mice mimics aspects of human MASLD and spontaneous tumorigenesis

As an exemplar of chronic stress, we studied a high-fat diet (HFD)-mediated liver injury model. In agreement with prior work, we found HFD C57BL/6 mice developed obesity, elevated serum cholesterol, increased alanine aminotransferase (ALT) levels, decreased serum albumin, and impaired glucose tolerance, indicative of hepatocellular damage, synthetic dysfunction, and diet-induced systemic insulin resistance (Fig. 1A-C, S1)^31^. Histologically, HFD livers exhibited (initially pericentral) steatosis, lobular inflammation, pericellular “chicken-wire” fibrosis, and hepatocellular ballooning (Fig. 1D-H, S1; see Supplementary Note 1 for additional contextualization against human disease progression). Spontaneous HCC developed without additional genotypic or chemical manipulation, which we validated in a second mouse cohort at a different institution (see Methods; 16/18 HFD mice vs. 4/19 CD mice developing liver tumors by 18 months; Fig. S1). These systemic and histologic findings mimic aspects of human MASLD progression: simple steatosis at 6 months, followed by certain features of inflammation, fibrosis, and spontaneous HCC tumorigenesis at 12 and 15 months.

**Figure 1:**
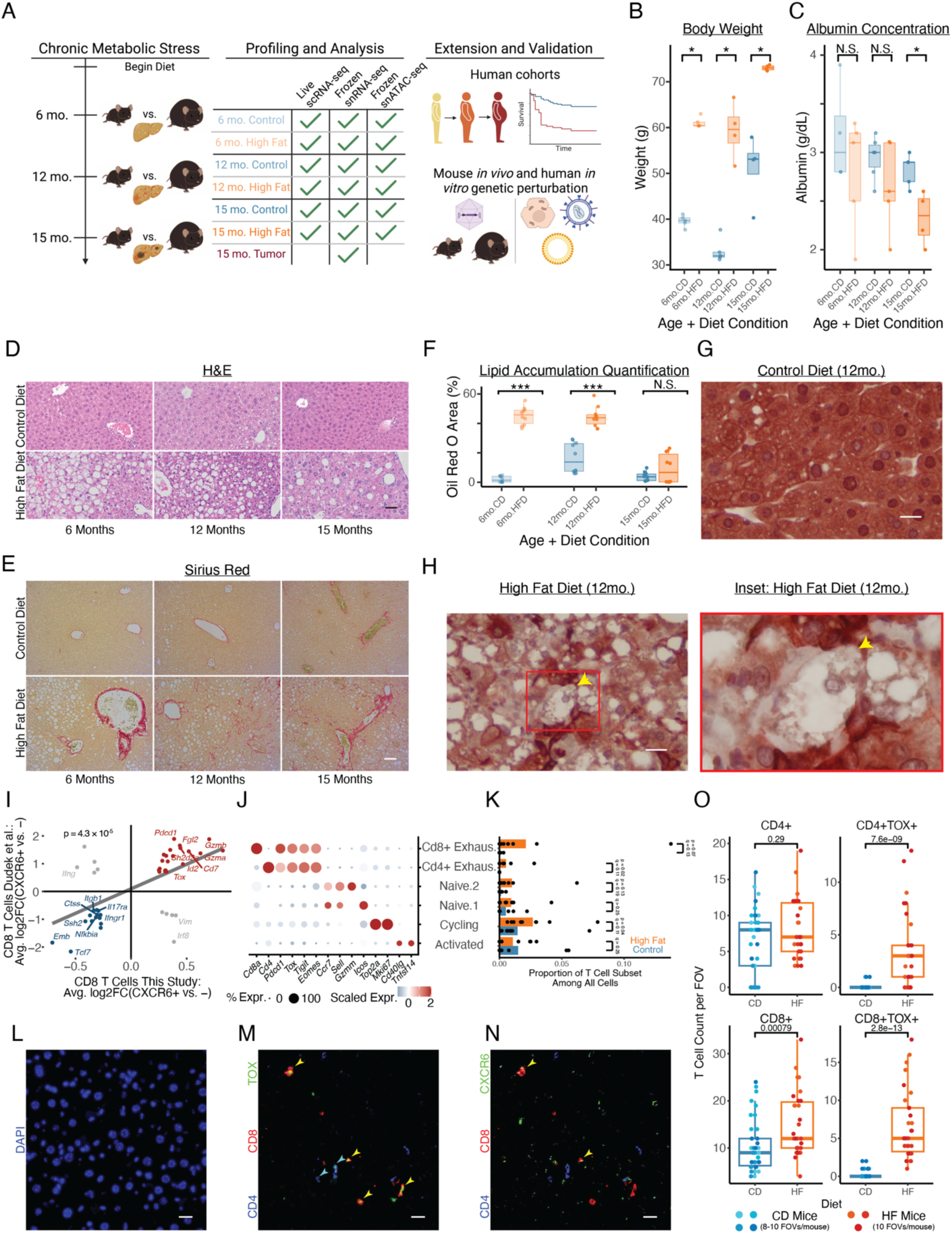
Diet-only mouse model of liver adaptations to chronic metabolic stress. (A) Study design schematic. (B) Mouse body weight. (C) Mouse blood albumin concentrations. (D-H) Histology of tissue morphology (D; scalebar=50µm), fibrosis (E; scalebar=50µm), lipid accumulation (F), and hepatocyte ballooning (CK8/18 staining; G-H; scalebar=15µm). (I) CXCR6^+^CD8^+^ T cell markers between human MASLD patients and our mouse model. (J) Annotated T cell subcluster markers (this study). (K) T cell composition with diet (this study). (L-N) Immunofluorescence of TOX^+^ and CXCR6^+^ T cells in mouse liver tissue (6mo HFD); (L) DAPI, (M) CD4/CD8/TOX, (N) CD4/CD8/CXCR6 (scalebar=50µm). (O) *In situ* T cell enrichment based on TOX status (6mo.HFD, this study). P-value in (I) calculated through Spearman’s correlation; all other p-values calculated with Mann-Whitney U test. * indicates p < 0.05; ** indicates p < 0.01; *** indicates p < 0.001.

To understand longitudinal adaptations to chronic metabolic stress, we conducted immune-biased live tissue single-cell RNA-seq (scRNA-seq; N = 17 mice, n = 23,819 cells), epithelial-biased frozen tissue single-nucleus RNA-seq (snRNA-seq; N = 18 mice, n = 79,408 cells), and frozen tissue snATAC-seq (N = 13 mice, n = 97,113 cells) for HFD and CD mice at each timepoint, recovering all major parenchymal, stromal, and immune subsets (Fig. S2-4). To further examine the fidelity of our MASLD murine model, we queried select non-parenchymal cell observations from the literature. For example, prior work has implicated human CXCR6^+^CD8^+^ T cells (expressing *TOX* and *PDCD1*) as drivers of autoaggressive hepatocyte killing in MASLD^32,33^. *Cxcr6*^+^*Cd8*^+^ T cells in our model recapitulated markers of these previously-described human cells, and T cells expressing exhaustion-related markers were compositionally enriched with HFD in our scRNA-seq dataset (Fig. 1I-K). Multiplexed immunofluorescence on mouse liver revealed that TOX^+^ T cells were strongly enriched *in situ* as early as 6 months (Fig. 1L-O). Likewise, prior work has identified human scar-associated macrophages (SAMac) as being both localized to cirrhotic regions and capable of activating collagen expression in stellate cells^34^. SAMac markers aligned with a cluster of macrophages in our scRNA-seq data; these SAMac-like mouse macrophages were enriched even at our earliest timepoint before overt fibrosis deposition (Fig. S5A-D). Collectively, these results indicate that our mouse model mirrors key molecular features of human MASLD.

### Metabolic stress drives tradeoffs between pro-survival pathways and core homeostatic functions

Given the diversity of homeostatic roles performed by hepatocytes, we focused on their dynamic stress responses and adaptations. Reasoning that genes with temporally-coordinated expression trajectories might capture linked biological processes, cellular responses, or regulatory targets, we defined four expression programs demonstrating progressive or rapid alteration in response to high-fat diet: 1) “Longitudinal Increase” and 2) “Sustained Upregulation”, which describe genes progressively or consistently elevated with long-term metabolic stress, respectively; and, 3) “Longitudinal Decrease” and 4) “Sustained Downregulation”, whose genes progressively decline or consistent lessening, respectively (Fig. 2A; Table S1).

**Figure 2:**
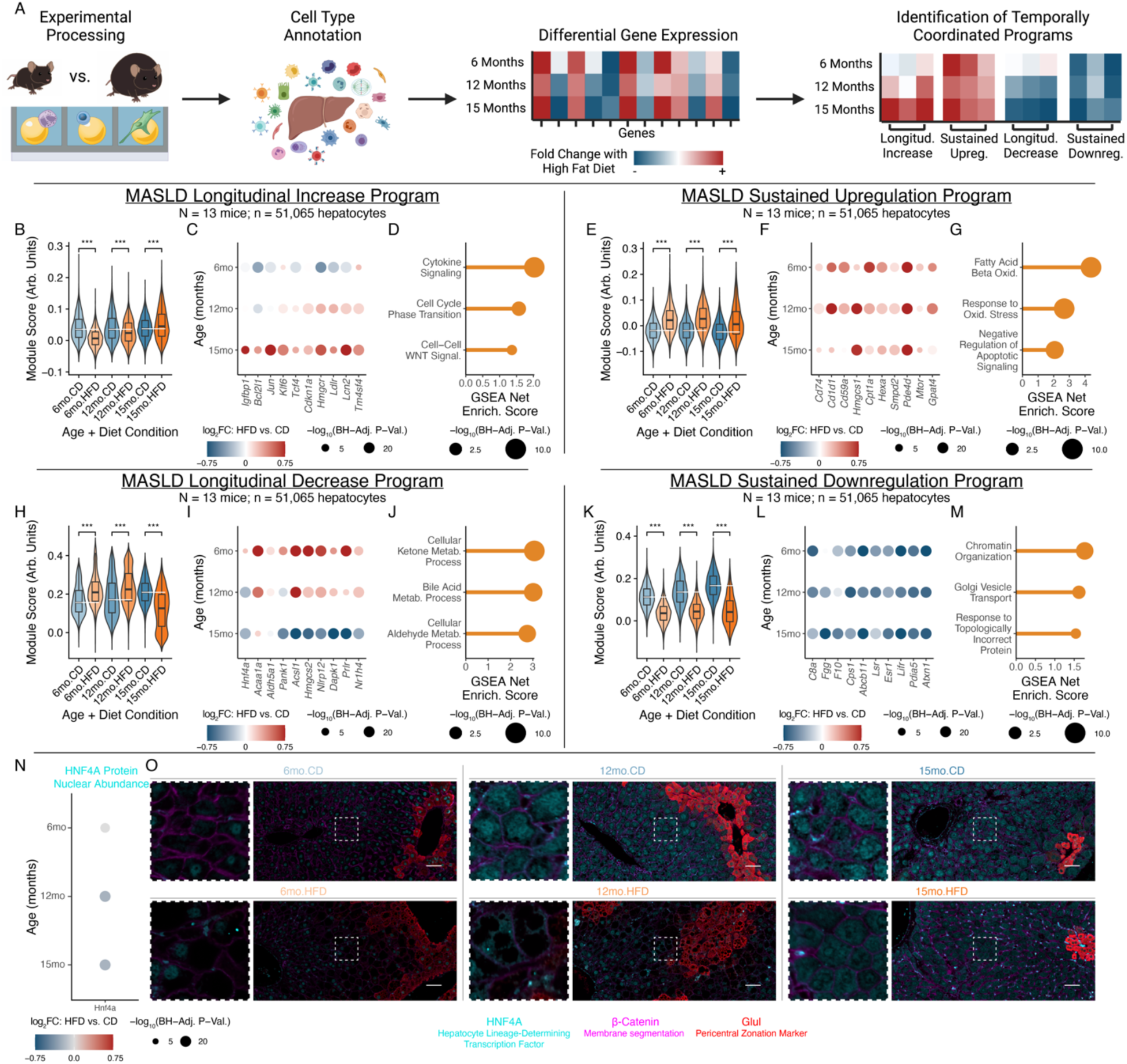
Dynamic adaptations of hepatocytes undergoing chronic metabolic stress. (A) Analysis schematic. (B-D) Longitudinal Increase program, with aggregate expression (B), expression changes of representative genes (C), or enriched GO:BP genesets (D). (E-G) Sustained Upregulation program, following (B-D). (H-J) Longitudinal Decrease program, following (B-D). (K-M) Sustained Downregulation program, following (B-D). (N) HNF4A protein nuclear abundance log_2_(fold-change) in hepatocytes with chronic metabolic stress, quantified through *in situ* tissue multiplexed immunofluorescence (N = 18 mice, 3 per age×diet condition; n = 350,842 nuclei). (O) Representative images of HNF4A protein nuclear abundance (scalebar=50µm). GSEA statistical testing through fgsea package; Mann-Whitney U-test used for all other tests; Benjamini-Hochberg correction applied for multiple testing. * indicates p < 0.05; ** indicates p < 0.01; *** indicates p < 0.001.

Examining the Longitudinal Increase program, long-term metabolic stress increased expression of genes involved in pro-survival pathways, WNT signaling, cholesterol metabolism, and intercellular signaling (Fig. 2B-D). For instance, *Igfbp1*, *Bcl2l1*, *Jun*, and *Klf6* have well-established roles in promoting hepatocyte survival and inhibiting apoptosis^35–38^. Increases in key WNT effectors (e.g., *Tcf4*) and regeneration-elevated regulators of cell cycle and senescence (e.g., *Cdkn1a*) suggest connections between stress responses and hepatocyte regeneration^39,40^. Linking stress responses and altered metabolism, hepatocytes also progressively upregulated the rate-limiting enzyme of cholesterol synthesis (*Hmgcr*) and key cholesterol uptake regulators (e.g., *Ldlr*), aligning with increases in cholesterol accumulation in hepatocytes^41,42^. Towards immune influences on hepatocyte phenotypes, we also observed upregulation of inflammation-responsive, regeneration-associated, and HCC-elevated genes like *Lcn2* (whose hepatocyte-specific knockout impairs acute regenerative capacity) and *Tm4sf4* (whose liver-specific overexpression increases inflammatory mediators like TNF and toxin-mediated acute liver damage)^43,44^. Globally, externally-defined gene sets related to cytokine signaling, cell cycle regulation, and WNT signaling were enriched among the Longitudinal Increase program (Fig. 2D; Table S2).

The Sustained Upregulation program, capturing aspects of the hepatocyte response to metabolic stress that are maintained over time, provided further examples of hepatocytes shifting to prioritize pro-survival responses under metabolic stress (Fig. 2E-G; Table S3). Hepatocytes increased expression of receptors mediating immune interactions and exerting pro-survival or anti-apoptotic effects, including *Cd74* (which promotes hepatocyte survival, in addition to antigen presentation, through binding to MIF)^45,46^, *Cd1d1* (which mitigates iNKT-mediated inflammation and hepatocyte apoptosis)^47^, and *Cd59a* (which prevents formation of complement membrane attack complexes on hepatocytes)^48^. Specific cholesterol synthesis and lipid oxidation enzymes were also upregulated, including *Hmgcs1* (catalyzing cytoplasmic acetoacetyl-CoA to HMG-CoA) and *Cpt1a* (rate-limiting enzyme for fatty acid beta-oxidation)^49,50^, as were metabolic regulators (e.g., *Mtor* and *Gpat4*) and enzymes catalyzing signal transduction metabolites (e.g., *Hexa*, *Smpd2*, *Pde4d*)^51–55^.

However, upregulation of pro-survival, WNT-associated, and immune interaction-related responses came at the expense of processes normally associated with the professional homeostatic responsibilities of hepatocytes (Fig. 2H-J; Table S4). *Hnf4a* was progressively downregulated with long-term metabolic stress; this is particularly notable given its role as a master regulator of hepatocyte identity, with wide-ranging effects on hepatocyte functions and fate specification during development^56^. Concordantly, the Longitudinal Decrease program also included enzymes related to peroxisomal oxidation (e.g., *Acaa1a*), aldehyde processing (e.g., *Aldh5a1*), CoA biosynthesis (*Pank1*; rate-limiting), and acyl-CoA processing (e.g., *Acsl1*)^57–61^. Progressive decreases in *Hmgcs2*, the rate-limiting enzyme of ketogenesis, were notable given that: 1) ketogenesis and cholesterol synthesis compete for the same starting metabolites^62^; and, 2) many cholesterol synthesis enzymes (including *Hmgcr*) followed opposing trajectories. Genes capable of driving cell death and serving as tumor suppressors, like *Nlrp12* and *Dapk1*^63,64^, were also downregulated. Regulators of hepatocyte phenotype, like *Prlr* and *Nr1h4* (encoding FXR), were likewise reduced with long-term metabolic stress. *Prlr* serves as a receptor for hormones including prolactin and growth hormone, and is upregulated by estrogens – notable given MASLD’s dramatically lower incidence in pre-menopause women^65,66^. The bile acid-activated nuclear receptor FXR, meanwhile, has been targeted in clinical trials via agonists including obeticholic acid, but the Phase III REVERSE trial recently failed to reach its primary endpoint^67^; we speculate that this may indicate opportunities for improved patient stratification based on expression and/or protein abundance of FXR.

Hepatocytes’ Sustained Downregulation program also reflected reductions in core hepatocyte and liver functions including secreted complement (*C8a*; also *C6*, *C8b*), coagulation factors (e.g., *Fgg*, *F10*), urea cycle (e.g., *Cps1*; rate-limiting enzyme), and bile acid regulation (e.g., *Abcb11*) (Fig. 2K-M; Table S5)^68–70^. *Lsr* regulates circulating triglyceride levels, and its experimental knockout has been shown to increase weight gain^71^, suggesting maladaptive repercussions of its diet-induced downregulation. We also observed downregulation of genes involved in proteostasis (e.g., *Pdia5*) and chromatin interactions (e.g., *Atxn1*)^72,73^. Suggestive of protection against hepatocyte self-destruction and complementing genes captured in other modules, downregulation of complement protein production (e.g., *C6*, *C8a*, *C8b*) harmonizes with the role of *Cd59a* (Sustained Upregulation program) in preventing complement-mediated hepatocyte death. *Esr1* encodes estrogen receptor alpha^74^, and its downregulation aligns with progressive losses of *Prlr* (Longitudinal Decrease program). Likewise, *Lifr* knockout is associated with increases in *Lcn2* secretion (Longitudinal Increase program) and elevated tumorigenesis^75^, potentially connecting metabolic stress-induced downregulation of an upstream receptor to subsequent pro-inflammatory, cancer-associated hepatocyte responses.

Towards validating dynamic shifts in hepatocyte phenotypes under chronic stress, we focused on HNF4A given its essential role in regulating hepatocyte differentiation and function. *In situ* mouse liver immunofluorescence demonstrated progressive decreases in HNF4A nuclear protein abundance with chronic metabolic stress, aligning with its Longitudinal Decrease transcriptional trajectory (Fig. 2N-O). We additionally cultured human liver HepG2 cells in lipid-rich media, and observed gene expression changes and functional metabolic and synthetic dysregulation concordant with sustained *in vivo* hepatocyte shifts (Fig. S5E-K, Methods)^76,77^

Thus, hepatocytes adapt to chronic metabolic stress by increasing expression of pro-survival responses, including anti-apoptotic effectors, WNT signaling, and intercellular interaction mediators. This pro-survival adaptation comes at the expense of genes linked to core hepatocyte identity and professional functions, including multiple classes of secreted molecules, transcription factors (including HNF4A and FXR), and metabolic enzymes.

### Adaptations to metabolic stress extend to human MASLD progression, exhibit extreme manifestations in cancer, and are prognostic for cancer survival

We next examined whether chronic stress adaptations in hepatocytes connected to tumor phenotypes, predicted long-term disease outcomes, and extended to human patient cohorts. We performed non-invasive MRI on mice fed a high-fat diet for 15 months to identify potential tumors, then dissected lesions that corresponded to grossly abnormal areas (Fig. 3A). Histologic evaluation supported liver tumors’ classification as HCC, with preserved liver architecture but also pleomorphism, prominent nucleoli, and hyperchromasia (Fig. 3B). Using snRNA-seq, we also found that tumors in our mouse model recapitulated human HCC markers, such as WNT and Notch pathway members (e.g., *Axin2*, *Tbx3*, *Ctnnb1, Lgr5*, *Jag1*), IGF signaling (e.g., *Igfbp1*, *Igf2r*), HCC diagnostic biomarkers (e.g., *Afp*, *Gpc3*), and proliferation (e.g., *Mki67*) (Fig. 3C, S6)^35,78–81^. More holistic analyses revealed enrichment for previously defined liver cancer marker sets with the expected directionality (Fig. S7A; Table S6-7)^82^.

**Figure 3:**
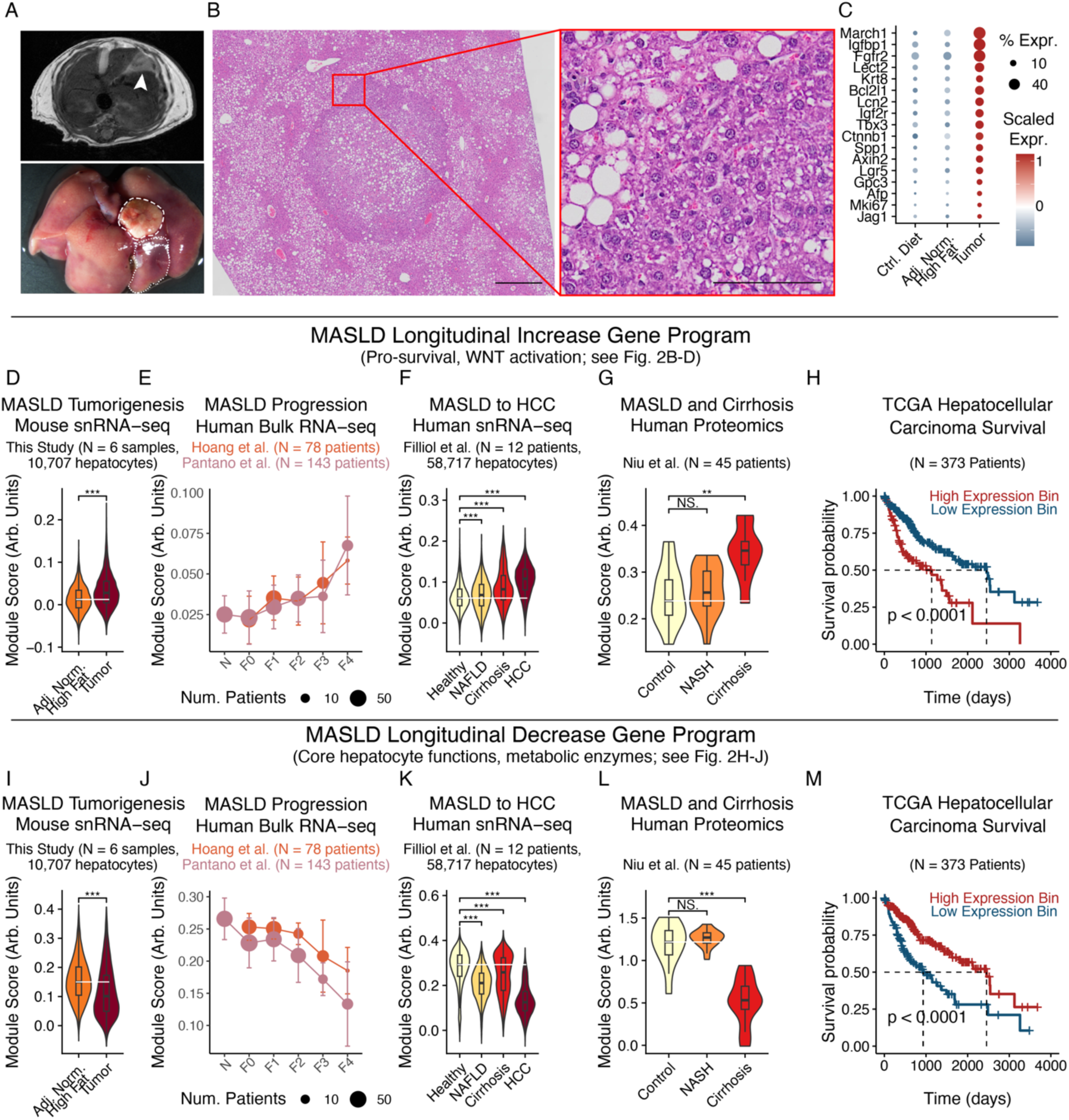
Chronic stress adaptations extend to human cohorts and connect to cancer phenotypes and outcomes. (A) MRI (top) and gross imaging (bottom) of mouse model spontaneous HCC. Dashed line indicates tumor; dotted indicates adjacent normal. (B) H&E of mouse model spontaneous HCC (left scalebar=500µm; inset scalebar=100µm). (C) HCC marker expression in mouse model spontaneous tumors. (D-H) Longitudinal Increase program expression in mouse snRNA-seq tumor cells and adjacent normal hepatocytes (D), human bulk liver RNA-seq (E), human snRNA-seq hepatocytes and tumor cells (F), human bulk liver proteomics (G), or human HCC survival outcomes (H). (I-M) Longitudinal Decrease program expression, following (D-H). Survival outcome p-values calculated with log-rank test; all other p-values calculated using Mann-Whitney U test with Benjamini-Hochberg correction. * indicates p < 0.05; ** indicates p < 0.01; *** indicates p < 0.001.

We next sought to understand whether and how chronic metabolic stress adaptations in non-transformed hepatocytes aligned with tumorigenesis-associated alterations. Tumor cells exhibited elevated expression of the Longitudinal Increase program relative to adjacent, non-transformed hepatocytes (Fig. 3D). External human cohorts exhibited a similar pattern, with the Longitudinal Increase program positively associated with human MASLD severity across microarray, bulk RNA-seq, and snRNA-seq readouts (Fig. 3E-F; Fig. S7B)^76,83–86^. Protein-level abundances of the Longitudinal Increase program likewise increased across MASLD progression in human liver tissue proteomics (Fig. 3G)^87^. Finally, we found that the Longitudinal Increase program was further heightened within human HCC and prognostically stratified HCC patient survival^88^, with high expression predicting worsened outcomes (Fig. 3F,H). Thus, the same pathways activated as adaptations to metabolic stress in non-transformed hepatocytes (e.g., pro-survival effectors, WNT activation) are additionally upregulated with tumorigenesis, so that tumor cells exhibit a heightened manifestation and extension of metabolic overload-induced adaptations.

The Longitudinal Decrease, Sustained Upregulation, and Sustained Downregulation programs also significantly associated with disease severity, tumor phenotypes, and patient survival outcomes (Fig. 3I-M; Fig. S7C-Q). As internal consistency, the Longitudinal Increase and Sustained Upregulation programs displayed opposite directionalities to the Longitudinal Decrease and Sustained Downregulation programs (i.e., worsened vs improved prognoses, respectively). Additionally, across studies, measurement modalities, and species, aggregate module scores were driven by similar underlying genes, with strong positive covariance between individual genes and overall module scores (Fig. S8A-D).

To further explore connections between early stress adaptations in non-transformed hepatocytes and later tumor phenotypes and outcomes, we investigated whether HCC-associated signatures were observed early in MASLD before overt tumorigenesis. Upon examining signatures of 1) cancer-linked chemical and genetic perturbations^82,89,90^; 2) HCC mutational subtypes^91^; and, 3) liver development and regeneration^92–96^, we found that hepatocytes take on gene expression patterns reminiscent of earlier developmental stages as well as the HCC S1 subclass even early in MASLD: development-associated expression programs (driven by genes including *Krt8*, *Sox9*, and *Cd24a*, Fig. S8E) were increased not only in HCC, but even at early disease stages in the MASLD progression in non-transformed hepatocytes across species and measurement modalities (Fig. 4A-E); likewise, marker genes of the human HCC S1 subclass, characterized by aberrant WNT activation, were elevated in hepatocytes across species at early MASLD stages and further elevated with tumorigenesis (Fig. 4F-J, Fig. S8F)^97^. Notably, β-catenin regulates WNT signaling activation and is a commonly mutated HCC driver gene^15,16^, suggesting pathway-level convergence between stress-induced transcriptional alterations and literature-established genomic drivers.

**Figure 4:**
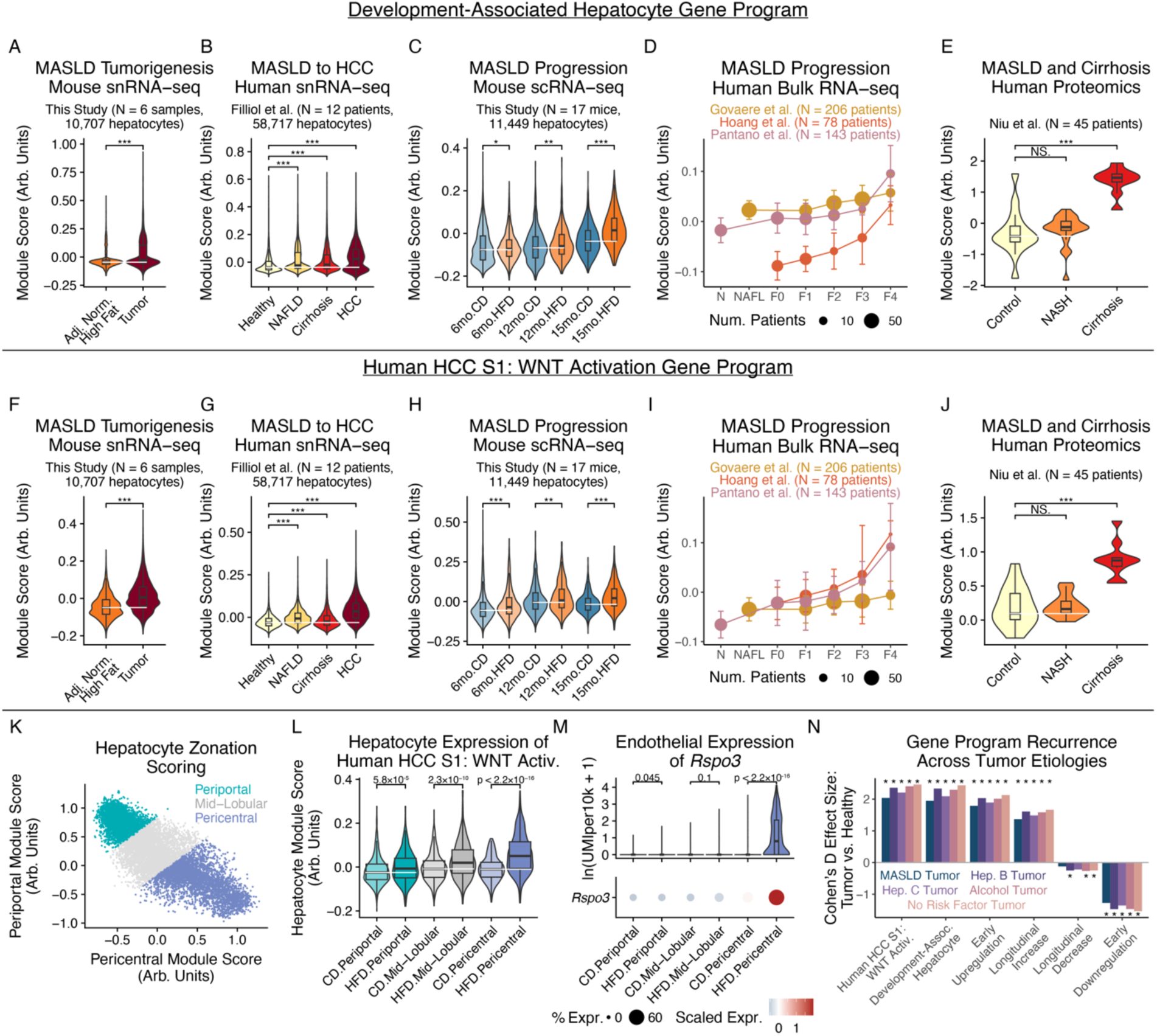
Chronic metabolic stress induces cancer signatures at early stages of MASLD progression. (A-E) Development-associated program expression in mouse snRNA-seq tumor cells and adjacent normal hepatocytes (A), human snRNA-seq hepatocytes and tumor cells (B), mouse snRNA-seq hepatocytes (C), human bulk liver RNA-seq (D), or human bulk liver proteomics (E). (F-J) Human HCC S1 WNT Activation program expression, following (A-E). (K) Mouse hepatocyte zonation scoring and annotation. (L) HCC S1 WNT Activation program expression by hepatocyte zonation and diet. (M) Endothelial *Rspo3* expression by zonation and diet. (N) Tumor-vs-healthy expression differences of this work’s hepatocyte adaptation programs, split by tumor etiology. All p-values calculated using Mann-Whitney U test with Benjamini-Hochberg correction. * indicates p < 0.05; ** indicates p < 0.01; *** indicates p < 0.001.

To better understand the broader spatial context of our observations, we inferred hepatocytes’ spatial positions via periportal-vs-pericentral markers (Fig. 4K)^98^. Pre-malignant pericentral hepatocytes exhibited larger increases in the WNT activation-associated HCC S1 signature than did periportal hepatocytes (Fig. 4L). Pericentral endothelial cells are a key source of secreted WNT ligands^99^. Upon similarly inferring endothelial cell zonation, we found that pericentral endothelial cells (but not periportal or mid-lobular endothelial cells) upregulate expression of *Rspo3*, *Wnt2*, and *Ctnnb1* with chronic metabolic stress, potentially aligning with spatially-structured signaling circuits (Fig. 4M, S9A-J; see Supplementary Note 2 for a discussion of the activation of hepatocyte chronic stress adaptation programs during acute regeneration and analyses of intercellular interactions potentially shaping hepatocyte adaptation programs). In examining whether these patterns were etiology-specific vs. generalizable, we found that hepatocytes’ metabolic adaptation programs were consistently dysregulated across HCC tumors linked to metabolic disorders (i.e., MASLD, alcohol-related liver disease), viral infection (i.e., hepatitis B and hepatitis C virus), or no known risk factors (Fig. 4N). Internally-consistent directionalities across HCC etiologies support the generalizability of these hepatocyte functional adaptations in wide-ranging disease microenvironments.

Overall, these results help to link metabolic stress-induced adaptations in non-transformed hepatocytes to later tumor phenotypes: extensions to human cohorts, similarities to later tumor states, and predictive power for disease severity and HCC patient survival. The directionality of effects on human HCC survival further helps propose interpretations for these axes of hepatocyte adaptation: while elevated processes could plausibly be linked to improved survival of individual cells, they incur longer-term repercussions through reductions in the expression of genes related to hepatocyte identity-defining features and professional functions, as well as early activation of pathways that may eventually contribute to tumorigenesis and worsened survival at later disease stages.

### Epigenetic dysregulation and WNT pathway priming under chronic metabolic stress

Having defined hepatocytes’ longitudinal adaptations to metabolic stress, we sought to obtain mechanistic insights into the epigenetic landscape and cell-intrinsic regulatory factors shaping hepatocyte expression patterns under stressful disease microenvironments (Fig. S4). Comparing hepatocyte metabolic adaptation programs across genomic regulatory layers, we found concordance between transcriptional expression and epigenetic accessibility (Fig. S10A).

To identify transcription factors (TFs) altered by metabolic stress in hepatocytes, we examined genome-wide accessibility of chromatin peaks containing TF binding motifs^100,101^ in our snATAC-seq dataset (Fig. 5A). We observed progressive increases in motif accessibility for members of the AP-1 complex (e.g., FOS, JUN), which play roles in responses to cellular stressors and mediate long-lasting epigenetic rewiring and tissue memory of inflammation in other compartments^23,102^. Increased TEAD motif accessibility suggests involvement of the Hippo pathway, which regulates liver regeneration and development^103,104^. Reinforcing our inference of early activation of WNT pathway and development-associated programs, we observed increased motif accessibility for WNT-associated TFs (e.g., TCF7, LEF1) and TFs active during liver development (e.g., SOX4, SOX9)^105,106^. As TFs with increased motif accessibility across both early and long-term metabolic stress, we found PPAR members (regulating liver metabolic pathways and serving as a leading MASLD drug target, but also driving stemness and regeneration phenotypes in intestinal stem cells), RXRA-associated nuclear receptor TF complexes, and NFE2L1 (a key regulator of cellular responses to oxidative stress)^107–111^.

**Figure 5:**
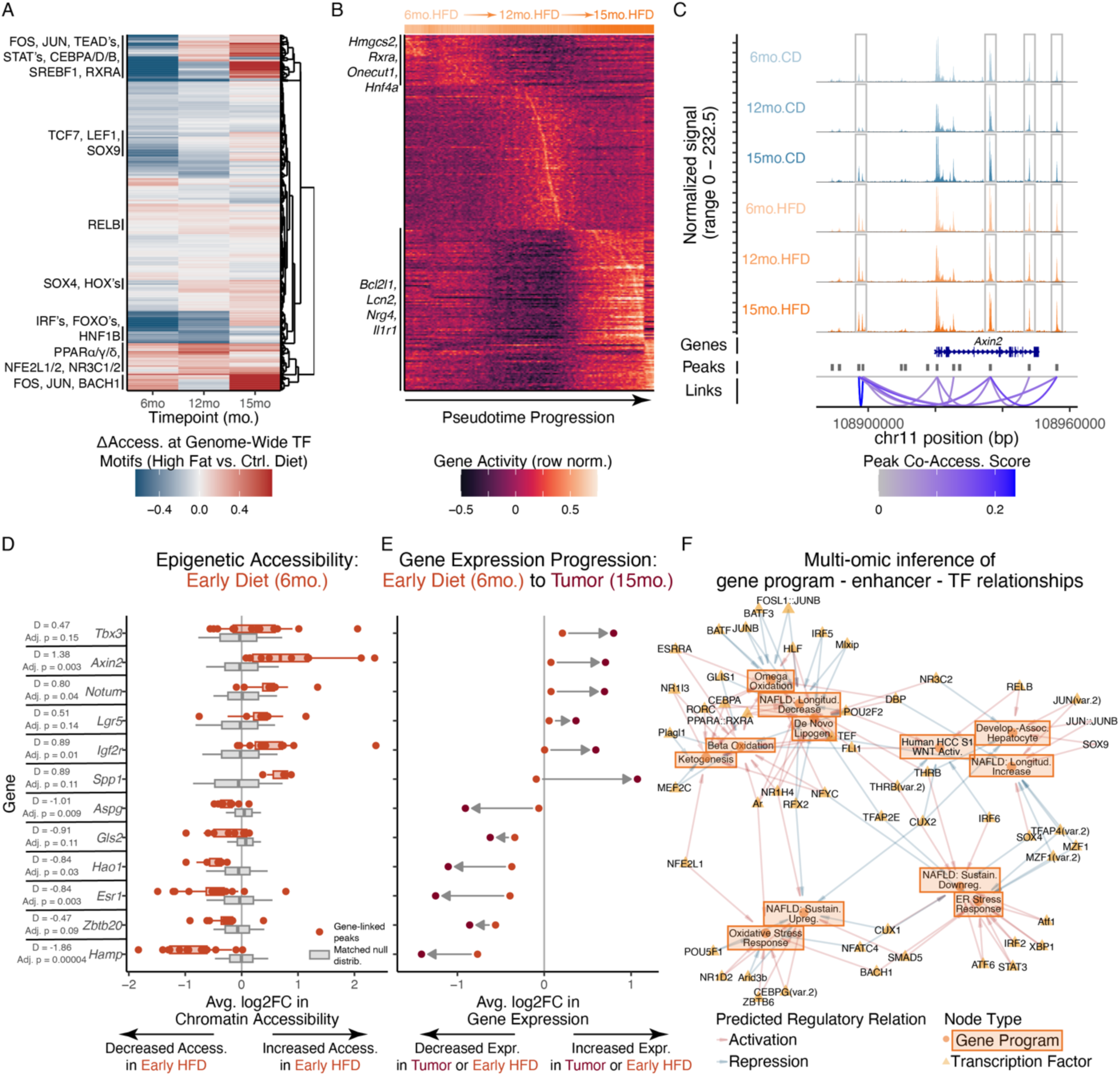
Chronic metabolic stress drives altered epigenetic trajectories and WNT pathway priming. (A) Chromatin accessibility deviation of TF motifs across age and diet conditions. (B) Epigenetic gene activity trajectories of HFD hepatocytes ordered by pseudotime progression. (C) Coverage plot of epigenetic accessibility at *Axin2* gene locus across age and diet. (D) Accessibility changes of gene-linked peaks at 6-month timepoint, between actual observed HFD-vs-CD fold-changes (red) and GC-and-background-matched null distribution (gray). (E) Expression changes of genes at 6-month HFD-vs-CD (red) or 15-month tumor-vs-adjacent-normal (brown) comparisons. (F) Computationally-inferred network of TFs predicted to regulate MASLD-relevant gene programs. All p-values calculated using Mann-Whitney U test with Benjamini-Hochberg correction; effect size quantified through Cohen’s D.

Towards higher-resolution insights into temporally-varying chromatin states, we conducted pseudotemporal analyses to order hepatocytes according to smooth gradients in chromatin accessibility, prioritizing gene loci dynamically altered by chronic metabolic stress (Fig. 5B, S10B-E). Chromatin trajectories reinforced our multi-omic, cross-species analyses of hepatocyte adaptations: members of the Longitudinal Decrease program including *Hmgcs2*, *Acsl1*, *Scp2*, *Hnf4a*, and *Rxra* exhibited maximal chromatin accessibility early in the high fat diet pseudotime progression, whereas members of the Longitudinal Increase program like *Lcn2*, *Bcl2l1*, and *Il1r1* peaked towards the pseudotime terminus. Progressive decreases in chromatin accessibility at *Hnf4a*’s gene locus are notable given its role as a master regulator of hepatocyte identity^112^, and aligns with its transcriptional and proteomic downregulation, decreases in professional hepatocyte functions, and increases in developmental marker expression. These (pseudo)temporal patterns were not observed in hepatocytes from control diet mice, supporting distinctive trajectories of stress-induced chromatin remodeling (Fig. S10B-F).

We additionally sought to identify genes where epigenetic alterations preceded and presaged transcriptomic shifts as these may indicate stress-induced epigenetic priming: chromatin remodeling establishing accessible epigenetic landscapes prior to transcriptional alterations, thereby priming cells for later activation and (dys)function^113^. We leveraged co-accessibility between intergenic chromatin peaks and promoters or gene bodies to create peak-gene linkages, capturing enhancer-gene regulatory interactions despite potentially large genomic distances^114^. The regeneration-upregulated, WNT/β-catenin target *Axin2* provides an example, with several distal chromatin regions co-accessible with *Axin2*’s promoter/gene body and elevated in accessibility with HFD across timepoints (Fig. 5C). More broadly, a variety of WNT-associated (e.g., *Tbx3*, *Axin2*, *Lgr5*, *Notum*) and HCC-linked (e.g., *Spp1*, *Igf2r*) genes exhibited increased epigenetic accessibility but only small changes in transcription at our earliest timepoint (6 months; Fig. 5D-E). However, these genes were strongly upregulated after tumorigenesis months later (15 months; Fig. 5E). Genes with the opposite directionality (i.e., early decreases in chromatin accessibility preceding more extreme transcriptional downregulation with longer-term metabolic stress) included: 1) metabolic enzymes and secreted proteins (e.g., *Aspg*, *Hao1*, *Hamp*); 2) suppressors of HCC and fetal hepatocyte-associated phenotypes (*Gls2* and *Zbtb20*, respectively); and, 3) *Esr1* as a receptor for MASLD-protective estrogen signaling (Fig. 5E,F)^115,116^.

Thus, paralleling hepatocytes’ transcriptomic and proteomic stress adaptations, discovery of early WNT- and HCC-linked chromatin changes that foreshadow transcriptional shifts suggests that even early exposure to metabolic stress may establish a permissive chromatin landscape that contributes to later activation of regeneration-, development-, and cancer-associated pathways.

### MATCHA prioritizes causal transcription factors shaping MASLD-associated phenotypes

Computational nomination of driving TFs for functionally-important gene programs (e.g., disease-linked stress adaptation programs in hepatocytes) remains an unsolved, open problem (see Supplementary Note 3 for contextualization of prior work). To accomplish this, we developed MATCHA (Multiomic Ascertainment of Transcriptional Causality via Hierarchical Association), a computational framework to map user-specified gene programs (e.g., arbitrary biological processes, disease (mal)adaptations, etc.) to distal enhancers and program-specific TF activities (see Methods). In brief, MATCHA links gene programs to cell-type-specific distal enhancers by identifying chromatin regions co-accessible with the gene program’s promoters or gene bodies. MATCHA then prioritizes causal regulators by determining TF motifs whose accessibility at program-coaccessible enhancers likewise covaries with program transcriptional expression. MATCHA further optionally incorporates: 1) concordance across datasets towards robust regulatory inference (e.g., across species, single-cell vs. bulk measurements, etc.); and, 2) identification of TFs co-regulating multiple gene programs. MATCHA therefore enables prioritization of TFs driving arbitrary gene programs while also modeling context-dependent functions via cell type- and tissue-specific gene regulatory landscapes (see Data and Materials Availability).

As proof-of-concept, we examined two metabolic-stress-relevant processes with known driver TFs: ER stress response and beta-oxidation (defined externally through GO:BP)^82,89,90^. MATCHA recovered ground-truth causal TFs for these external test cases: the top two TFs prioritized for GO:BP ER stress response genes were XBP1 and ATF6 (i.e., two of three well-established master regulatory TFs) (Fig. S11A-G)^117^. The top 5 TFs for GO:BP beta-oxidation, meanwhile, included FXR/*NR1H4* and PPARA (whose agonists have advanced to Phase III clinical trials for MASLD)^110,118^, and NR1I2, CEBPA, and NR1I3 (all with preclinical evidence for regulation of liver metabolism and lipid accumulation; Fig. S11H-N)^119,120^.

To identify core TFs mediating hepatocyte longitudinal stress adaptations and early induction of development-associated and HCC-linked cell states, we applied MATCHA to construct a bipartite network of regulatory relationships between gene programs and TFs, supported by epigenetic and transcriptional evidence (Fig. 5F). Our network successfully recapitulated known ground truths, but also nominated targets for experimental validation. In addition to previously-discussed well-known drivers of ER stress (i.e., XBP1, ATF6) and beta-oxidation (i.e., PPARA, NR1H4/FXR), our network captured hepatocyte developmental regulation (i.e., SOX9 driving development states), hormonal influences on metabolism (i.e., androgen receptor driving beta-oxidation^121,122^), and stress response mediators (i.e., NFE2L1/NRF1 driving oxidative stress response^123^). We also note inferred regulatory links between: 1) THRB motifs; 2) activation of hepatocyte identity-linked Longitudinal Decrease and Sustained Downregulation programs; and, 3) repression of development-associated and HCC S1 WNT programs. We note that the THRB agonist resmetirom is currently undergoing the Phase III MAESTRO-NASH trial^124^. We chose 15 TFs (20 isoforms) to prioritize for experimental validation (Fig. S11O).

### RELB and SOX4 mediate wide-ranging MASLD adaptation and tumor-associated functional phenotypes

To validate MATCHA-nominated drivers of hepatocyte metabolic adaptation, we conducted arrayed human *in vitro* genetic perturbations^125^. We created HepG2 cells stably overexpressing TF isoforms and cultured them in lipid-rich media, followed by: 1) scRNA-seq to validate TF effects on hepatocyte transcriptomic phenotypes (n = 10,522 cells; median 417 cells per TF isoform); and, 2) live-cell imaging and immunofluorescence to validate TF effects on functional metabolic and cancer-associated phenotypes (Fig. 6A, S12; Table S8-10). Transcriptomic profiles of negative controls (non-transduced and BFP-transduced cells) clustered with each other by media condition but not transduction status, suggesting: 1) specificity of measured responses (via separation from TF-transduced cells); and, 2) preserved responses to lipid-induced metabolic stress following transduction (Fig. 6B). As a positive control, we confirmed that overexpression of PPARA in lipid-rich culture led to upregulation of known targets, including *HMGCS2*, *CPT1A*, *PLIN2*, *G6PC1*, and *PCK1* (Fig. 6B, S12C)^126^.

**Figure 6:**
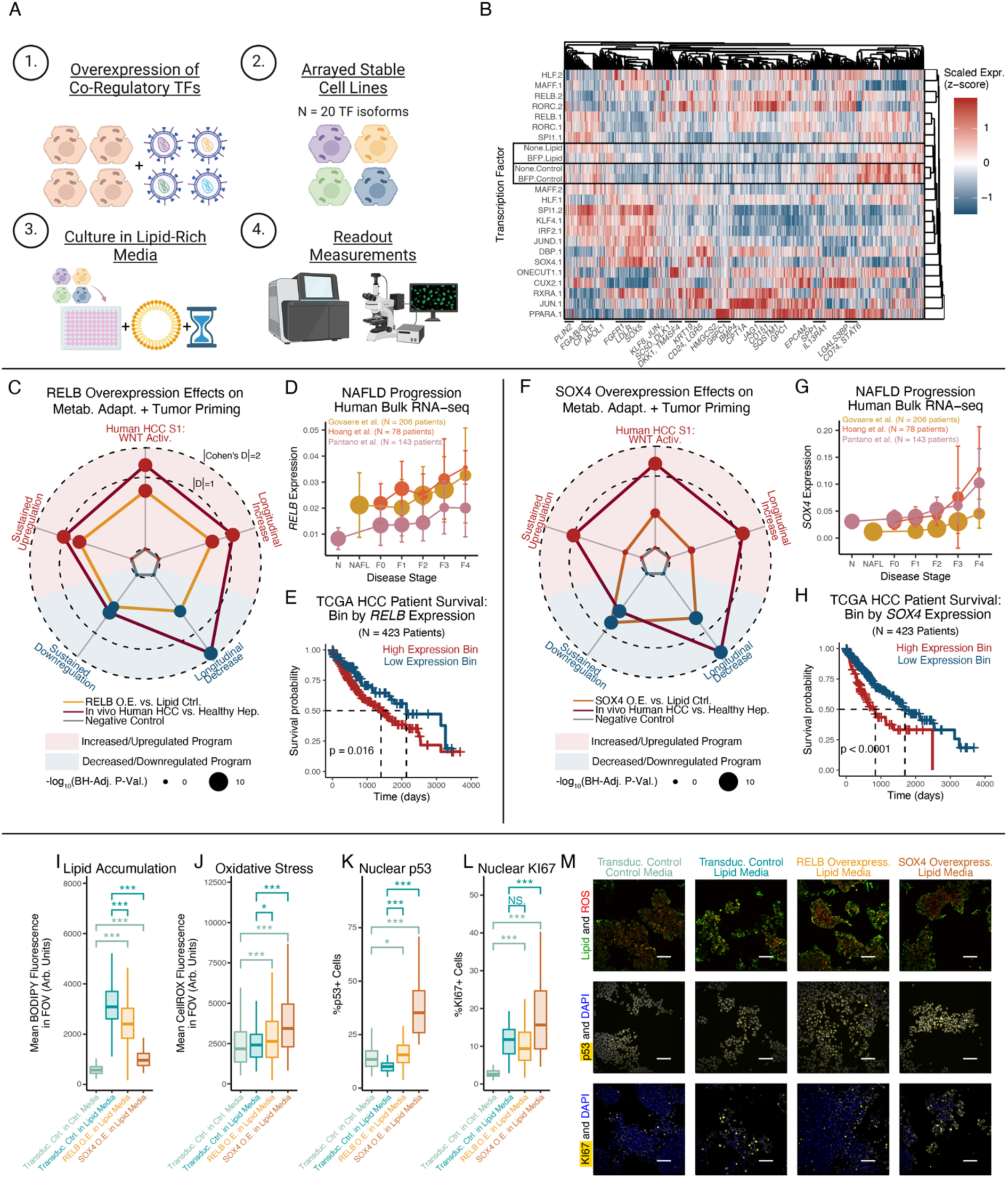
Human *in vitro* validation of RELB and SOX4 as regulators of hepatocyte metabolic (mal)adaptation. (A) Experimental design schematic. (B) Pseudobulked TF expression profiles. Where applicable, “.1” and “.2” indicate isoforms of the same TF gene. (C) RELB’s effect sizes on gene program expression. Radius equals |Cohen’s D| if concordant directionality with human MASLD/HCC, and 0 if discordant. (D) RELB expression across human MASLD progression. (E) HCC human survival stratified RELB by expression. (F-H) Transcriptomic regulatory effects of SOX4, following (C-E). (I-L) Lipid accumulation (I; BODIPY 493), ROS accumulation (J; CellROX), nuclear p53 (K), and nuclear KI67 (L). (M) Representative microscopy images supporting (I-L) (scalebar=100µm). Survival outcome p-values calculated with log-rank test; all other p-values calculated using Mann-Whitney U test with Benjamini-Hochberg correction. * indicates p < 0.05; ** indicates p < 0.01; *** indicates p < 0.001.

With successful recovery of known TF-target relationships, we evaluated how MATCHA-nominated TFs regulated hepatocyte stress adaptations. Downstream targets of RELB, RORC, and MAFF overexpression included developmental markers (e.g., *EPCAM*), pro-survival effectors (e.g., *KLF6*, *JUN*), cholesterol synthesis and fatty acid oxidation enzymes (e.g., *SC5D*, *CPT1A*), and receptors involved in HCC-relevant immune interactions (e.g., *CD74*, *LGALS3BP*)^46,127^. Overexpression of SOX4 and ONECUT1 (active during liver maturation) drove downstream WNT pathway targets and regulators (e.g., *DKK1*, *DLK1*) and developmental markers (e.g., *CD24*, *KRT19*), among others^105,128^. Additionally, SOX4 (along with JUND, KLF4, and CUX2) upregulated metabolism-associated genes, such as lipid droplet-associated *PLIN2* and *CPT1A* (rate-limiting enzyme for beta-oxidation)^21,129^. With individual MATCHA-nominated TFs capable of regulating wide-ranging aspects of hepatocytes’ stress adaptations, we proceeded to focus on RELB and SOX4 given the strength of their effect on metabolic adaptation-associated transcriptional and functional states.

Overexpression of RELB (a member of the non-canonical NF-κB signaling complex) in lipid-rich media drove transcriptional shifts consistent with hepatocytes’ *in vivo* long-term metabolic adaptations: elevation of the Longitudinal Increase, Sustained Upregulation, and HCC S1 signature programs, and decreases in the Longitudinal Decrease and Sustained Downregulation programs (Fig. 6C). Specific RELB targets included increases in development-associated and regeneration-linked markers (e.g., *CD24*, *CDKN1A*, trend towards *LGR5*), decreases in hepatocyte secreted protein products (e.g., *FGB*, *FABP1*), and upregulation of HCC-associated genes (e.g., p62/*SQSTM1*, *CD151*, *LGALS1, CD74*) (Fig. S13A). Contextualizing against *in vivo* shifts, we observed that RELB drove effect sizes that were smaller than, but of comparable magnitude to, tumor-vs-healthy differences (Fig. 6C), suggesting both the breadth and strength of RELB on hepatocyte stress adaptations. Towards *in vivo* MASLD relevance, RELB expression increased with fibrosis stage across multiple human cohorts, and RELB as a single marker stratified patient survival, associating with worsened outcomes (Fig. 6D-E). For further human *in vivo* evidence of RELB regulation of hepatocytes’ stress adaptations, we examined how human HCC patients’ copy number variations (CNVs) at the RELB locus altered expression of downstream gene programs, analogous to a “natural genetic perturbation experiment” in human tumors^130^. RELB CNVs drove significant changes in *RELB* transcription and predicted: 1) significant increases in the development-associated, HCC S1 WNT Activation, and Sustained Upregulation programs; 2) significant decreases in the Sustained Downregulation and Longitudinal Decrease programs; and, 3) non-significant elevation of the Longitudinal Increase program (Fig. S13B-H). Thus, human *in vitro* genetic perturbations and human *in vivo* HCC CNVs support RELB as a driver of hepatocyte stress adaptation programs, loss of cell identity, and development-associated and cancer-linked states.

Overexpression of SOX4 (SRY-box transcription factor 4, active during liver development) in lipid-rich media also depleted genes associated with hepatocyte cellular identity and function including TFs (e.g., *NR1H4*, *PPARA*, trend towards *HNF4A*), metabolic enzymes (e.g., *GPD1*, *AKR1D1*, *DPYD*, *EHHADH*), and secreted protein products (e.g., *SERPINF2*, *PLG*, *FABP1*, *FGG*) (Fig. 6F, S13I). In contrast, SOX4 increased expression of developmental markers (e.g., *CD24*, WNT targets *LGR5* and *NKD1*), as well as genes with functional effects in HCC (e.g., *CD151*, *MDK*, *AIFM2*, *AKR1C2*, *ROBO1*). As a broader, orthogonal examination of SOX4 driving loss of hepatocyte identity, we examined how SOX4 overexpression altered genes enriched in hepatocytes relative to all other cell types^131^, as a proxy for the distinguishing features and cell identity of hepatocytes. 89% of statistically-significant hepatocyte identity-associated genes were downregulated by SOX4 expression, supporting SOX4 as driving loss of hepatocyte identity (Fig. S13J). Towards human *in vivo* relevance, we found that *SOX4* expression increased with MASLD severity and stratified HCC patient survival, associating with worsened survival (Fig. 6G-H).

Functionally, overexpression of RELB and SOX4 decreased cellular lipid accumulation, but also caused accompanying dose-dependent increases in ROS accumulation (Fig. 6I-J, M). These effects may be mediated by shared downstream target genes linked to reduced lipid accumulation (e.g., increased *ABCC1* and *CPT1A*, decreased *FABP1*) and elevated oxidative stress (e.g., decreased *GSTA1*, decreased *MT1E* and *MT2A*) (Fig. S13I)^132–137^. TF-mediated tradeoffs between lipid and ROS accumulation may align with prior work proposing a potentially protective role for lipid droplets, which can sequester lipid species that might otherwise drive lipotoxicity or increase ROS levels upon processing and oxidation^138,139^. To connect these factors to longer-term disease outcomes, we further examined the effect of RELB and SOX4 on proliferation. Overexpression of RELB or SOX4 each increased nuclear accumulation of p53 protein (Fig. 6K,M). SOX4 overexpression drove increased proliferation as measured by nuclear KI67; RELB overexpression did not drive significant changes in nuclear KI67, which could be interpreted as preserved proliferative capacity despite cellular stress (Fig. 6L-M; see Supplementary Note 4 for discussion of p53 regulation of hepatocyte phenotypes and proliferation, and especially connections to metabolism, development-associated states, WNT, and AP-1 signaling).

Through human *in vitro* genetic perturbation experiments and extensions to human cohorts, we validated RELB and SOX4 as causal regulators of stress-induced transcriptional and functional tradeoffs between hepatocyte identity and dysfunction-associated phenotypes, unifying hepatocytes’ metabolic adaptations around specific regulatory nodes.

### HMGCS2 is a metabolic mediator of hepatocytes’ adaptation and shifts towards cancer-associated phenotypes

We finally sought to demonstrate the *in vivo* importance of dynamic shifts in hepatocyte stress adaptation programs. We chose to focus on HMGCS2 (3-hydroxy-3-methylglutaryl-CoA synthase 2) and the regulatory effects of ketogenesis-cholesterol metabolic rewiring, given HMGCS2’s: 1) role as the rate-limiting enzyme of ketogenesis; 2) strong upregulation with PPARA activation (Fig. S12C); 3) cross-species expression decreases with longitudinal metabolic stress; and, 4) association with low expression and worsened HCC survival (Fig. S14A-D). We generated a mouse model with hepatocyte-specific knockout of HMGCS2, analogous to *Hmgcs2* expression decreases with the natural stress adaptation progression (*Hmgcs2*^fl/fl^; *Alb-Cre*) (Fig. S14A-D). After 6 months on either HFD or CD, steatosis occurred in mice on HFD with either wildtype (WT) or hepatocyte-specific HMGCS2 knockout (HepKO), but was comparatively less severe in CD mice of either genotype (Fig. 7A). Liver damage (measured by circulating transaminases) and cholesterol increased with the combination of HFD and HMGCS2 HepKO (Fig. S14E-F). Validating the HMGCS2 knockout, HMGCS2 abundance was decreased throughout the liver lobule via immunohistochemistry, and we observed reduced circulating ketone bodies following a 24-hour fast (Fig. 7B-C). However, HMGCS2 HepKO did not alter circulating glucose concentrations or HFD-driven weight gains, indicating specificity of effects on ketogenesis (Fig. 7D-E).

**Figure 7:**
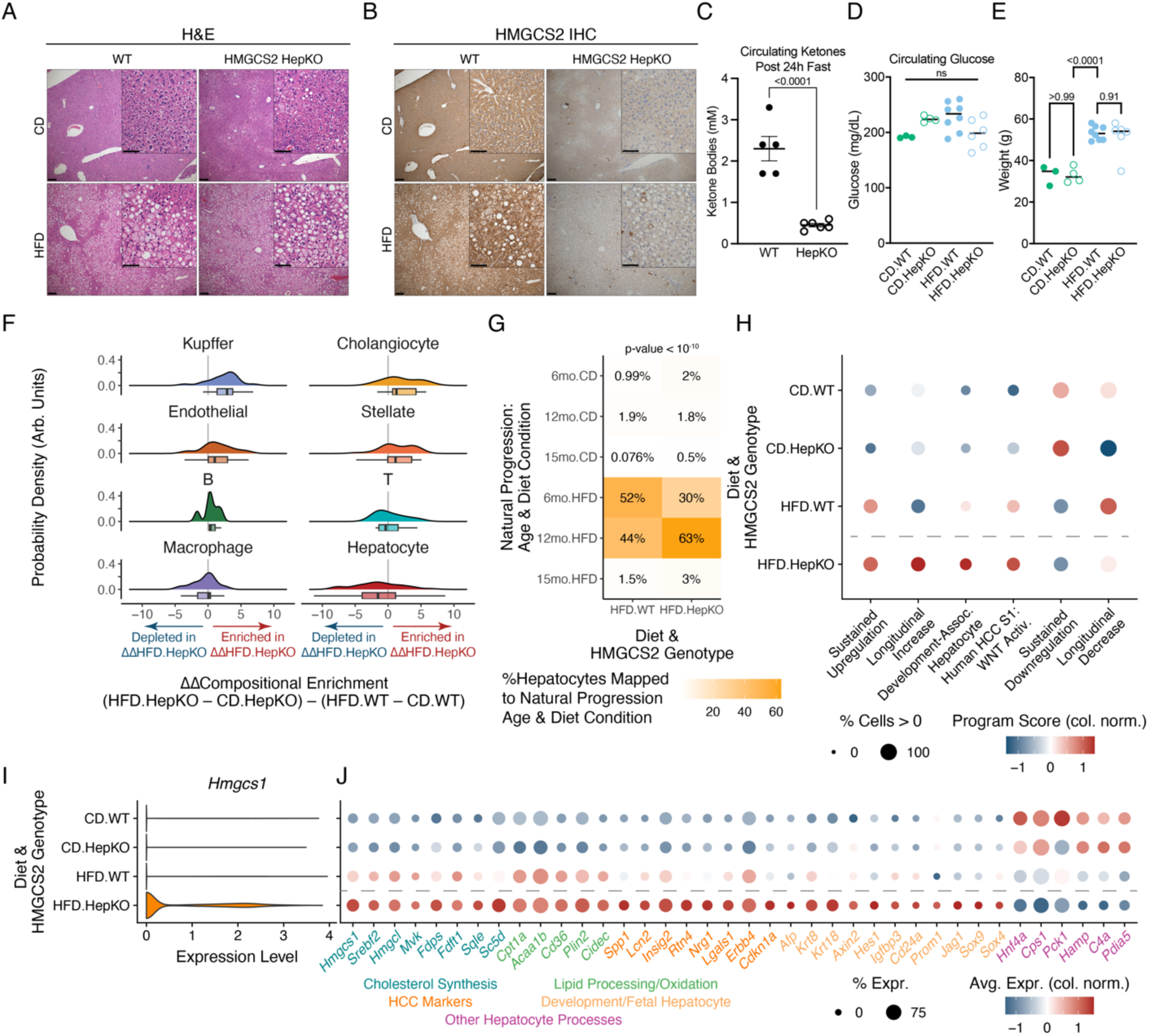
*In vivo* validation of HMGCS2 as a regulator of hepatocyte metabolic (mal)adaptation. (A) H&E staining of wildtype (WT) or *Hmgcs2*^fl/fl^; *Alb-Cre* (HMGCS2 HepKO) mice on CD or HFD (main image scalebar=200µm; inset scalebar=100µm). (B) HMGCS2 immunohistochemistry. (C) Circulating ketone body concentrations after 24-hour fast. (D-E) Circulating glucose concentrations (D) or body weights (E). (F) HMGCS2 HepKO and diet-induced compositional shifts via Milo-derived neighborhoods. (G) Reference mapping of hepatocytes from HMGCS2 HepKO cohort mice to this study’s natural stress adaptation progression. (H) Gene program expression with diet and HMGCS2 genotype. (I) Expression of *Hmgcs1* with diet and HMGCS2 genotype. (J) Gene expression across diet and HMGCS2 genotype. P-value in (G) calculated using Fisher’s exact test; all other p-values calculated using Student’s t-test.

To investigate the effects of HMGCS2 loss on liver adaptation under chronic metabolic stress, we conducted snRNA-seq at the 6-month timepoint across HMGCS2 genotypes and diet conditions (N = 8 mice, n = 27,119 cells) (Fig. S15; Table S11). Neighborhood-based analyses of our snRNA-seq data indicated compositional shifts specific to the metabolic stress response of HepKO mice, but not that of WT mice (Fig. 7F). To validate our compositional analyses, we focused on Kupffer cells, where immunohistochemistry against the Kupffer cell-predominant marker CD68 supported their compositional enrichment with the interaction of metabolic stress and hepatocyte-specific HMGCS2 loss (Fig. S14G-I).

To understand the causal role of HMGCS2 and ketogenesis on hepatocyte phenotypes, we mapped gene expression profiles of HFD.WT and HFD.HepKO hepatocytes to the natural progression of chronic metabolic stress (Fig. 7G). Approximately equal proportions of HFD.WT hepatocytes mapped to the natural progression 12-month (44%) and 6-month (52%) conditions. However, with HFD.HepKO, over twice as many hepatocytes mapped to the 12-month HFD condition (63%) as to the 6-month HFD condition (30%), indicative that HMGCS2 loss causes accelerated dysfunctional phenotypes associated with later stages of hepatocyte stress adaptation.

More directly, we examined how HMGCS2 HepKO under metabolic stress altered hepatocyte stress adaptation programs. HMGCS2 HepKO on HFD led to extreme expression states, even relative to diet-induced shifts in WT mice: larger elevation of Sustained Upregulation, Longitudinal Increase, Development-Associated, and HCC S1 WNT Activation programs, but also further reductions of Sustained Downregulation and Longitudinal Decrease programs (Fig. 7H). To uncover specific targets, we prioritized genes exhibiting emergent transcriptional shifts with metabolic stress and HMGCS2 HepKO. Diverse cholesterol synthesis-related genes were upregulated with the combination of HMGCS2 HepKO and HFD (e.g., *Hmgcs1*, *Srebf2*, *Hmgcl*, *Mvk*), in line with compensation at the metabolic branch point between ketogenesis and cholesterol synthesis (Fig. 7I,J). Additionally, HMGCS2 HepKO on HFD induced cell states directly associated with long-term dysfunction, including HCC-linked intercellular signaling proteins (e.g., *Spp1*, *Lcn2*, *Lgals1*) and development-associated markers (e.g., *Afp*, *Axin2*, *Hes1*, *Cd24a*, *Sox9*). Loss of HMGCS2 on the background of HFD also accentuated decreases in expression associated with hepatocyte identity (e.g., *Hnf4a*) and function (e.g., *Cps1*, *Pck1*, *Hamp*, *C4a*, *Pdia5*). Finally, connecting HMGCS2 to MATCHA TF inference and experimental validation, HMGCS2 loss under chronic metabolic stress significantly upregulated *Sox4*.

Thus, *in vivo* genetic perturbation validated wide-ranging regulatory effects of HMGCS2 itself, but also more broadly the functional importance and interpretation of our stress adaptation programs. Premature HMGCS2 loss induced accelerated damage and extreme manifestations of transcriptional programs, including compensatory cholesterol synthesis, HCC phenotypes, development-associated states, and loss of hepatocyte canonical functionality. Experimental *in vivo* modeling of accelerated stress adaptation progression (through genetic manipulation) further supported adaptation programs derived in this work as fundamental, functionally-important axes of hepatocyte response to environmental stressors.

## Discussion

During chronic stress, cells must balance survival against performing their professional functions. We investigated how the liver manages longitudinal tradeoffs under environmental stressors through the paradigm of chronic metabolic overload, which precipitates progressive steatosis, inflammation, fibrosis, cirrhosis, and malignant transformation. In addition to its pressing clinical need, the biological context of MASLD offers opportunities to understand how initial cellular adaptations connect to longer-term tissue dysfunction, disease pathogenesis, and patient prognosis.

We developed a diet-only mouse model that exhibits functional, histologic, and cellular phenotypes paralleling human MASLD. Our longitudinal single-cell multi-omics datasets, ranging from early steatosis to late spontaneous tumorigenesis (with matched diet controls) along with harmonized human MASLD/HCC transcriptomic and proteomic cohorts, provide rich computational resources on the intersection of aging and metabolic stress. Importantly, as our mouse model does not require genetic manipulation or exogenous chemical insults, it may be a broadly-translatable experimental resource for further investigations. We acknowledge that HFD mice do not develop significant bridging fibrosis or nodule formation, possibly due to lifespan differences between mice and humans, thereby creating opportunities for further model development for investigations focused on stellate cells and fibrosis.

We demonstrated that chronic metabolic stress induces development-linked and cancer-associated adaptations in hepatocytes to the detriment of cellular identity and professional tissue functions. Hepatocytes increased genes related to: 1) early developmental stages; 2) pro-survival and anti-apoptotic effectors; and, 3) intercellular signaling (including WNT). In contrast, hepatocytes downregulated genes underpinning homeostatic roles, including diverse secreted proteins and metabolic enzymes. These findings were corroborated across multiple human cohorts, and recapitulated across epigenetic, transcriptomic, and proteomic readouts.

Suggesting discovery of generalizable stress adaptation responses, gene programs uncovered in this work exhibited extreme manifestations in acute regeneration and across HCC risk factors (consistent across etiologies including MASLD, alcohol, and viral infections). These connections support further investigations of uncovered gene programs’ generalizability as core, conserved axes of hepatocyte adaptation to diverse stressors. Such conserved axes in cross-disease cohorts could uncover broadly applicable vs. disease-specific therapeutic vulnerabilities and patient stratification hierarchies. As potential mechanisms for conserved cross-disease stress adaptation programs, distinct molecular changes associated with each etiology may converge on similar intracellular mediators (e.g., analogous to diverse PAMPS or DAMPs converging on TLR activation)^140^. Alternatively, each etiology may drive recurrent microenvironmental or immune signals that in turn produce consistent hepatocyte phenotypes (e.g., regeneration contributions of both pro-inflammatory IL-6 and pro-fibrotic TGFB1)^141,142^.

Progressive decreases in lineage-determining HNF4A, decreases in genes mediating canonical hepatocyte functions, and increases in developmental markers raise the question of how stress adaptations overlap or contrast with dedifferentiation. Partial hepatocyte dedifferentiation occurs following acute stressors like experimentally-induced partial hepatectomy and acetaminophen overdose, where subsets of hepatocytes downregulate canonical functions and assume fetal-like phenotypes to enable regeneration and restoration of liver mass^4–6,99,143^. Genetically-induced priming of dedifferentiation improves hepatocyte survival during subsequent acute stressors, but long-term dedifferentiated states predispose to liver failure and worsened survival in HCC^92,144–146^. In cancer, dedifferentiation is acknowledged as a recurrent hallmark, where cancer cells unlock fetal, plastic cell states associated with elevated proliferation and tumor progression^147,148^. One potential model to unify our findings involves clinically-relevant chronic stresses driving partial dedifferentiation-associated states even in non-transformed hepatocytes. In this model, the trajectory of hepatocytes’ progressive stress adaptations would increasingly draw upon the liver’s regenerative capacity for improved survival of individual cells, but with deleterious repercussions: 1) maladaptive tradeoffs with the liver’s tissue-level functions and homeostatic setpoints; and, 2) early induction and priming of similar transcriptional programs as occur with tumorigenesis and worsened cancer outcomes. Future demonstrations functionally linking stress adaptation and de-differentiation (especially early in disease progression in non-transformed hepatocytes) may further delineate transcriptional and epigenetic contributions to liver failure and elevated cancer incidence.

Extending hepatocytes’ immediate adaptations, we also found WNT signaling members and HCC markers exhibited increased chromatin accessibility months prior to transcriptional upregulation with tumorigenesis. These findings suggest stress adaptations incurring not just direct changes, but also epigenetic priming for later activation of cancer-associated states. In other organs and diseases, AP-1 synergizes with context-specific TFs to maintain poised epigenetic landscapes and memory of past inflammation^23^; future work could investigate the minimal requirements and timescales needed for epigenetic dysregulation and eventual phenotypic manifestations. For instance, in the skin and pancreas, inflammatory bouts lasting only a handful of days are sufficient to drive effects upon triggers administered months later (e.g., improved wound healing, increased cancer risk)^19,20,24,149^, which would suggest even short stressors can instill persistent tissue memory and altered tissue function. In contrast, in the nasal epithelium, modulation of cytokine signaling can partially revert dysfunctional progenitor memory of allergic memory, towards therapeutic restoration of aspects of homeostatic baseline^102,150^. Hepatocytes’ primed accessibility at AP-1 binding motifs and WNT-related loci occurred at relatively early disease stages which precede when many patients exhibit symptoms^7^. Thus, future investigations into the timescales of stress adaptations’ initiation and persistence are motivated by not only fundamental biological understanding but also the longer-term clinical relevance of whether hepatocyte tissue memory and lasting maladaptive phenotypes may be seeded prior to disease diagnosis. With weight loss capable of stabilizing or even reversing histologic features of MASLD, whether and to what degree hepatocytes’ memory of chronic stress is reversible or establishes lasting (mal)adaptive responses even after return to normal weights could shape long-term implications of ongoing trials and novel therapeutic avenues for MASLD^111,151,152^.

To extend from discovering axes of cellular stress adaptations to uncovering their causal regulators, we developed MATCHA, a computational framework to prioritize TFs driving arbitrary, user-specified gene programs. MATCHA infers gene program – co-accessible enhancer – causal TF triads by leveraging: 1) multimodal -omics data across layers of genome regulation, temporal trajectories, and species; and, 2) context-dependent regulatory relationships ensuring output predictions capture disease- and cell type-specific TF activity. When applied to externally-defined biological processes, MATCHA recovered their well-established ground truth drivers. When applied to uncover “central hub” TFs with strong co-regulatory connections across multiple stress adaptation programs derived in this work, MATCHA re-discovered targets of multiple Phase III clinical trials, but also highlighted comparatively less-explored TFs that we experimentally validated. A strength of MATCHA is its emphasis on incorporating tissue and cell type-specific data as the basis for computational inference of regulatory relationships, capturing context dependence of chromatin landscapes and TF activity. Future improvements of MATCHA could include prioritizing sets of TFs needed to drive the breadth of a gene program (e.g., beyond top TFs prioritized in isolation that may have overlapping, noncomprehensive target genes). Alternatively, in addition to predicting TF regulatory effects on a gene program, we could seek to elucidate differential gene regulatory network structures for each TF under different contexts (e.g., longitudinal stress adaptation stages) to reveal context-specific alterations in enhancer binding and downstream target genes.

Validating MATCHA predictions, RELB and SOX4 indeed regulated hepatocytes’ stress adaptation transcriptional programs and functional metabolic phenotypes in human *in vitro* genetic perturbation experiments. RELB constitutes the transcriptional effector of the non-canonical NF-κB signaling pathway^153^. In the liver, significant prior work investigated cytoplasmic regulation of NF-κB kinase complexes, as well as canonical NF-κB signaling’s roles in tumorigenesis and hepatocyte survival under inflammatory conditions^26,154–160^. However, canonical and non-canonical NF-κB differ significantly in terms of input signaling mechanisms and output phenotypic effects: they are activated by distinct ligands, involve separate intracellular interactions, and culminate in different TFs^153,161,162^. Thus, the role of RELB in hepatocytes’ progressive remodeling under metabolic stress has received less attention^163^. Our work points towards novel contributions of RELB to driving hepatocytes’ (mal)adaptations to chronic stress, across disease associations, transcriptional targets, and functional effects. Likewise, SOX4 is active during fetal liver development and epithelial progenitor fate specification towards cholangiocytes^93,105^. In HCC, SOX4 predicts worsened survival, and ectopic Sox4 overexpression *in vivo* decreases hepatocyte identity features and drives metaplasia^164–166^. However, SOX4’s involvement in adult hepatocytes’ metabolic regulation and adaptations has been less explored. Our work supports SOX4’s involvement in mature hepatocytes’ stress responses, with especially strong links to downregulation of canonical hepatocyte functions that align with the posited de-differentiation interpretation of hepatocytes’ stress adaptations. These results additionally support that even individual regulatory nodes can be sufficient to co-regulate and couple phenotypes with opposing functional associations and temporal trajectories during progressive stress adaptations.

While our human *in vitro* model allowed us to validate the cell-intrinsic effects of different TFs, future work could define their upstream activating signals (e.g., through co-culture or organoid systems enabling dissection of intercellular interactions, biochemical cues, mechanical microenvironments, etc.)^3,167–173^. Our intercellular signaling analyses nominated ligands associated with both MASLD severity and activation of RELB or SOX4 (Supplementary Note 2): LTβ activates both canonical and non-canonical NF-κB signaling, supports successful acute regeneration and mouse survival after partial hepatectomy, and was predicted to drive hepatocytes’ Sustained Upregulation program via production by compositionally-enriched T cells^159,161,174,175^. Likewise, TGFB1 activates SOX4^176^ and was predicted to drive hepatocytes’ Longitudinal Increase and development-associated programs via production from compositionally-enriched macrophages. SOX4 can additionally be activated by WNT signaling^177,178^, whose activation was supported at epigenetic, transcriptomic, and proteomic levels.

Towards demonstrating beneficial-vs-maladaptative repercussions of dynamic shifts in stress adaptation programs, our analyses highlighted opposing temporal trajectories of ketogenesis and cholesterol synthesis. These pathways compete for the same starting precursor metabolite of acetoacetyl-CoA (product of fatty acid oxidation)^62^. Recent work demonstrated that acetoacetyl-CoA accumulation may increase liver tumorigenesis risk via histone acetylation modifications that facilitate accessible, permissive chromatin landscapes^179^. However, as hepatocytes allocate flux through these metabolic pathways for processing acetoacetyl-CoA, their downstream products have opposing associations with hepatocyte health: free cholesterol can directly cause lipotoxicity^8^, while hepatocytes lack the necessary enzyme to catabolize ketone bodies and therefore export them to other organs^180^. Demonstrating maladaptive effects of ketogenesis decreases during naturally-occurring stress adaptations, metabolically-stressed *Hmgcs2*^-/-^ hepatocytes exhibited accelerated shifts towards phenotypes characteristic of later stages of chronic stress exposure: 1) compensatory increases in cholesterol synthesis enzymes; 2) extreme manifestations of adaptation gene programs; 3) upregulation of HCC markers and development-associated genes including *Sox4*; and 4) downregulation of lineage-determining *Hnf4a* and canonical hepatocyte functions. We previously showed that ketone bodies regulate intestinal stem cell regenerative capacity via epigenetic interactions^181^; future work could further elucidate precise molecular mechanisms and interactions by which ketogenesis, cholesterol synthesis, and associated metabolites regulate the hepatocyte response to metabolic stress.

One final important question is whether stress adaptation programs uncovered in this work are necessarily co-regulated or can be disentangled for selective modulation. Pareto analyses have investigated core archetypes of cellular contributions to collective functions, with hepatocytes accomplishing tissue-scale tasks through division of labor along the periportal-pericentral axis^182–184^. However, these analyses focused on steady-state healthy hepatocytes and did not consider dynamic axes of disease progression. Thus, disease contexts present additional complexity (but also opportunities) for understanding how cells and tissues navigate the state space of potential phenotypes and maintain essential functions despite external stressors. Our work sought to demonstrate the existence and functional implications of chronic stress adaptation programs by focusing on validation of transcriptional (RELB, SOX4) and metabolic (HMGCS2) mediators of wide-ranging dysfunction. However, future work could attempt to decouple these programs, therapeutically activating only beneficial features while mitigating otherwise-linked deleterious phenotypes. For instance, engineered transcriptional activators may enable support of both cellular survival and tissue function without priming of phenotypes associated with worsened cancer outcomes. Alternatively, targeting key enzymes to tune relative metabolic fluxes could provide greater control of tissue homeostatic setpoints, towards buffering of healthy function against environmental stressors.

In this work, we demonstrated how long-term stress drives adaptations that balance immediate cellular survival, homeostatic tissue functions, and long-term dysfunction. Through dissection of hepatocytes’ temporal adaptation trajectories, computational methods development to nominate cell-extrinsic and cell-intrinsic drivers, and experimental validation via human *in vitro* and mouse *in vivo* genetic perturbations, we coalesced diverse axes of hepatocyte adaptation around their specific causal factors. Ultimately, our work provides a foundation for revealing the principles behind cellular and tissue decision-making during stress, translating complex descriptions of disease dysfunction into unifying core mechanisms, and deriving fundamental connections of how even early stress can precipitate cellular adaptations and tradeoffs which lead to long-term dysfunction.

## Materials and Methods

### Mouse Husbandry and High Fat Diet

C57BL/6 mice in the MIT cohort were housed and cared for in accordance with the American Association for the Accreditation of Laboratory Animal Care and approved by MIT’s Committee on Animal Care. A long-term high-fat diet containing 60% kcal from fats (Research Diets D12492) was provided ad libitum to male mice starting at the age of 8-12 weeks for 6-15 months. Sex- and age-matched control mice were provided a purified Control diet containing 10% kcal fats and matched sucrose (Research Diets D12450J). C57BL/6 mice in the second mouse cohort at Brigham and Women’s Hospital were housed and cared for in accordance with the American Association for the Accreditation of Laboratory Animal Care and approved by Brigham and Women’s Hospital Committee on Animal Care, with diets started at the age of 4 weeks in these mice (Research Diets D12492 and D12450J). *Alb-Cre* mice (#003574) were purchased from Jackson Laboratory and described previously^185^. *Hmgcs2*^fl/fl^ were generated in-house and described previously^181^. The following mice were bred in-house: *Hmgcs2*^fl/fl^; *Alb-Cre*. Studies involving hepatocyte-specific loss of HMGCS2 were performed using littermates, whereby Cre-negative mice served as controls. Live animal imaging was arranged through the Koch Institute at MIT Animal Imaging and Preclinical Testing Core and performed using a Varian 7T MRI imaging system.

### Mouse Functional and Histological Characterization

- Cholesterol, ALT, Albumin: For cholesterol, ALT, and albumin measurements, blood was collected through cheek bleed or cardiac puncture in Microvette 200 Z-Gel containers (Sarstedt 20.1291). Briefly, samples were centrifuged immediately after collection, and serum was frozen at −80C before being submitted for IDEXX lab testing as coordinated by the Division of Comparative Medicine at MIT.
- HMGCS2 and CD68 IHC: Tissue was fixed in 10% normal buffered formalin prior to paraffin-embedding. 4-5 micron sections underwent deparaffinization and rehydration prior to antigen retrieval using Borg Decloacker RTU solution (Biocare Medical, BD1000G1) and a pressurized Decloaking Chamber (Biocare Medical, NxGen). Antibodies and respective dilutions used for immunohistochemistry are as follows: rabbit monoclonal anti-HMGCS2 (1:2000, Abcam, ab137043) and rabbit monoclonal anti-CD68 (1:500, Cell Signaling Technology, #97778) with dilutions performed in Signalstain Antibody Diluent (Cell Signaling Technology, #8112). Biotin conjugated secondary donkey anti-rabbit antibodies (1:500, Jackson ImmunoResearch) were used prior to Vectastain Elite ABC immunoperoxidase detection kit (Vector Laboratories, PK6100). Visualization was performed using Signalstain DAB substrate kit (Cell Signaling Technology, #8049). Counterstaining was performed with 50% Gill No. 1 hematoxylin solution (Sigma-Aldrich, GHS116). Images were acquired using an Olympus BX43 microscope with 4x and 10x objectives (Olympus UPlanSApo). Aperio Digital Slide Scanning at 20x magnification was performed for CD68 slides prior to analysis and quantification using QuPath software.
- Histologic Evaluation: H&E stained sections of liver tissue from 6, 12, and 15-month diet cohorts as well as identified tumor samples were reviewed in a blinded-manner by Dr. Vikram Deshpande and Dr. Ömer Yilmaz, both of whom serve as clinical hepatopathologists in the Department of Pathology at MGH.
- Sirius Red: For collagen staining, 5µm paraffin-deparaffinized sections were hematoxylin stained, washed in running tap water, and then stained with Picro Sirius Red for 1 hour (0.1% Direct Red 80 in picric acid).
- Oil Red O: 10µm frozen sections were briefly fixed, rinsed in isopropranol, and incubated in Oil Red O for 15min with hematoxylin counterstain.
- Glucose tolerance: To measure systemic insulin resistance, intraperitoneal glucose tolerance testing was performed in 6-hours fasted mice as described^186^.
- IHC (CK8/18): Staining was performed as described^187^ with the following modifications. Antigen retrieval was in buffer TE (pH=9) in a pressure cooker (Biocare Medical Decloaking Chamber, DC2002). A primary antibody already targeting K8/18 was used instead of mixing K8 & K18 1:1 (Progen, GP11; 1:400), detected with ImmPACT AMEC Red per manufacturer instructions (Vector Laboratories, SK-4285), and counterstained with hematoxylin.

### Multiplexed Immunofluorescence

Iterative, multiplexed immunofluorescence was conducted as previously described^188^ with the following modifications. Briefly, samples were deparaffinized and antigen retrieved in TE (pH=9) in a pressure cooker (Biocare Medical Decloaking Chamber, DC2002). The following antibodies were used: anti-HNF4A (Abcam, ab201460; 1:400), anti-CXCR6 (Thermo Fisher, PA5-79117; 1:125), anti-TOX (Abcam, ab237009; 1:50), anti-CD4 (Abcam, ab183685; 1:500), and anti-CD8A (Abcam, ab217344; 1:500). Imaging was conducted on a confocal Nikon Ti2 microscope with a Yokogawa CSU-W1. Primaries were incubated overnight at 4C in a humidified slide box. For quantitative measurement of nuclear localized HNF4A, Cellpose^189^ was used to segment nuclei based on DAPI signal with mean targeted diameter of 9µm (Cellpose optimized for hepatocyte nuclei). Nuclear masks from Cellpose were imported into CellProfiler^190^ along with HNF4a channels. Nuclei with area < 20μm^2^ were discarded to enrich for hepatocyte nuclei, and then mean fluorescence of nuclear segmentations in the HNF4A channel were computed using the MeasureObjectIntensity module.

### Live-Tissue Dissociation for scRNA-seq

Two-step collagenase liver perfusion was performed as previously described^99^. 15,000 cells were loaded onto a Seq-Well array for experimental processing. Samples were sequenced on an Illumina NextSeq 500/500, NextSeq 2000, or NovaSeq 6000.

### Frozen-Tissue Nuclei Isolation for Tandem snRNA-seq and snATAC-seq

For each sample, the following solutions and volumes were prepared for isolation of nuclei for tandem Seq-Well snRNA-seq and 10x v1.1 snATAC-seq. All solutions were pre-chilled and kept on ice except when actively handling the tissue or nuclei solution, and then immediately returned to ice.

- Base buffer: 100µL of 1M Tris-HCl pH 7.4, 20µL of 5M NaCl, 30µL of 1M MgCl_2_, 1000µL of 10% bovine serum albumin
- Wash buffer + RNase inhibitor: 230µL of base buffer, 20µL of 10% Tween-20, 1.7mL of nuclease-free water, 50µL of Sigma-Aldrich Protector
- Wash buffer + digitonin: 230µL of base buffer, 20µL of 10% Tween-20, 1.746mL of nuclease-free water, 4µL of 5% digitonin
- 1X lysis buffer: 230µL of base buffer, 20µL of 10% Tween-20, 20µL of 10% Nonidet P40 substitute, 1.736mL nuclear-free water
- Lysis dilution buffer: 230µL of base buffer, 1.77mL of nuclease-free water
- 0.1X lysis buffer: 200µL of 1X lysis buffer, 1.75mL of lysis dilution buffer, 50µL of Sigma-Aldrich Protector
- Diluted nuclei buffer: 48.75µL of 20X Nuclei Buffer (from 10x v1.1 scATAC-seq reagent kit), 926.25µL of nuclease-free water, 25µL of Sigma-Aldrich Protector
- PBS + 1% BSA + RNase inhibitor: 875µL PBS, 100µL of 10% BSA, 25µL of Sigma-Aldrich Protector

Flash-frozen pieces of liver tissue were kept on dry ice; if needed, a smaller piece of tissue (∼2-3mm diameter) was cut using a scalpel on a petri dish on dry ice. The tissue piece was placed into a Miltenyi C tube containing 2 mL of 0.1X lysis buffer, then homogenized using 2 iterations of the m_spleen_01 program on the gentleMACS tissue dissociator. Half of the solution was passed through a 40µm filter pre-wet with 1mL wash buffer + RNase inhibitor, and the other half of the solution was passed through a separate 40µm filter prewet with 1mL wash buffer + digitonin. The C tube was washed with 1mL of wash buffer + RNase inhibitor to capture remnant nuclei stuck to the tube side or lid, and 500µL was transferred to each separate tube. Each solution was transferred to a separate 15mL Falcon tube and centrifuged for 10min at 500g and 4^oC^, with brake set to 5 (out of a maximum of 10). The supernatant was aspirated, and each pellet was resuspended in 100µL of diluted nuclei buffer. Two 35µm filters were pre-wet with diluted nuclei buffer, and flow-through from the pre-wetting solution was removed to leave the tubes empty; each nuclei solution was then passed through the separate pre-wet 35µm filters using a P200 pipette. Each set of nuclei was counted using a hemocytometer, followed by snRNA-seq and snATAC-seq as previously described:

- 15,000 nuclei in 200µL of PBS + 1% BSA + RNase inhibitor as input to Seq-Well S^3^, as described in Hughes*, Wadsworth II*, Gierahn*, et al., *Immunity* (2020)^191^
- 7,000*1.53 nuclei in 5µL of diluted nuclei buffer as input to 10x snATAC-seq, as described in Chromium Next GEM Single Cell ATAC Reagent Kits v1.1 User Guide, CG000209 Rev F

A similar protocol was followed for nuclei isolation from: 1) the frozen tumor and adjacent normal tissue cohort; and, 2) the HMGCS2 HepKO cohort. The following modifications were made:

- As snATAC-seq was not conducted, the splitting of nuclei into a tube containing wash buffer + digitonin and subsequent parallel processing steps were omitted; all nuclei were passed through a single 40um filter pre-wet with 1mL wash buffer + RNase inhibitor.
- At the 35µm filter step, instead of diluted nuclei buffer, the filter was pre-wet with PBS + 1% BSA + RNase inhibitor solution.

Samples were sequenced on an Illumina NextSeq 500/500, NextSeq 2000, or NovaSeq 6000.

### Lentiviral Production

Plasmids encoding TF ORFs were obtained as generated in Joung et al., *Cell* (2023) (also deposited in Addgene MORF Collection).

600,000 Lenti-X 293T cells were plated in 2mL of media (DMEM + 10% FBS + 1% penicillin-streptomycin) in a 6-well plate and incubated overnight at 37^oC^ and 5% CO_2_ (Day 1). The following afternoon (Day 2), 1µg of psPAX2, 0.33µg of pMD2.G, 1.33µg of ORF plasmid, 5.3µL of P3000 reagent, and 125µL of Opti-MEM were mixed, added to a solution of 6.7µL of L3000 and 125µL of Opti-MEM (not mixed), and incubated for 15min at room temperature. The mixture was added to the Lenti-X 293T cells and incubated overnight. The following morning (Day 3), Lenti-X 293T media was replaced. The following afternoon (Day 4), Lenti-X 293T media was harvested and replaced; the overnight media was passed through a 0.45µm low protein binding filter, combined with Lenti-X Concentrator at a 3:1 media:Lenti-X ratio, and stored overnight at 4^oC^. The following afternoon (Day 5), Lenti-X 293T media was again harvested and added to the previous day’s media and Lenti-X Concentrator solution at 4^oC^ for 30min. The media was spun at 1500g for 45min at 4^oC^. Supernatant was aspirated, and the pellet was resuspended in 160µL for storage at −80^oC^ in 20µL aliquots.

### Lentiviral Transduction

For each TF ORF, 500,000 HepG2 cells were plated in 2mL of media (Advanced DMEM/F12 + 10% FBS + 1% penicillin-streptomycin) in a 6-well plate and incubated overnight at 37^oC^ and 5% CO_2_ (Day 1). The following morning (Day 2), lentiviral stocks of each TF ORF and polybrene were thawed to room temperature. Cells’ media was replaced with a 10µg/mL solution of polybrene in 2mL of HepG2 media, followed by addition of 8µL of lentivirus to separate HepG2 wells (i.e., arrayed format; 1 TF ORF per well). Cells were incubated with lentivirus overnight, and media was replaced the following morning (Day 3). The following afternoon (Day 4), media was replaced with a 1ug/mL solution of puromycin in HepG2 media.

To produce stable, arrayed HepG2 lines overexpressing each TF, cells were maintained in puromycin-containing media to select for successful transduction, with puromycin-containing media being changed every 2-3 days. Upon expanding to reach confluency in a 6-well plate, each TF ORF-overexpressing HepG2 line was passaged and replated in a 10cm dish. Upon reaching confluency in a 10cm dish, each TF ORF-overexpressing HepG2 line was passaged to freeze down cell stocks (1,000,000 cells in 1mL of 90% FBS + 10% DMSO at −80oC in a Mr. Frosty Freezing Container), followed by subsequent functional and transcriptomic assays in the metabolic stress of lipid-rich media.

### Liver Cell Lipid Culture, scRNA-seq, Functional Imaging, and Immunofluorescence

HepG2 cells overexpressing each TF ORF were seeded in a 96-well plate at a density of 4,000 cells/well. As lipid-rich, metabolically-stressful media, TF ORF-overexpressing cells were cultured in Advanced DMEM/F12 media containing 10% FBS, 500µM palmitic acid, 100µM oleic acid, and 1% penicillin-streptomycin. As controls, BFP-transduced HepG2 cells and non-transduced HepG2 cells were each cultured in lipid-rich media or control media (Advanced DMEM/F12 media containing 10% FBS, 600mM BSA control, and 1% penicillin-streptomycin). Each well received 200µL of its respective media condition (i.e., TF ORF-overexpressing HepG2’s in lipid-rich puromycin-containing media; BFP-transduced HepG2’s in lipid-rich or control puromycin-containing media; non-transduced HepG2’s in lipid-rich or control puromycin-free media). Media was changed 2 and 4 days after seeding, with assays occurring 7 days after seeding.

For scRNA-seq, HepG2 cells for each condition were seeded across triplicate wells. 7 days after seeding, cells were passaged, and triplicate wells for each condition were pooled and used as input for scRNA-seq using Seq-Well S^3^.

For functional imaging, cells were stained with BODIPY (2µM final concentration), CellROX Deep Red (1:500 final dilution), and Hoechst 33342 (5µg/mL final concentration) in PBS at 37^oC^ and 5% CO_2_. After 30min incubation, cells were washed 3 times with PBS, followed by imaging on an Opera Phenix at 37^oC^ and 5% CO_2_.

After functional imaging, cells were fixed in a solution of 4% paraformaldehyde in PBS for 10 minutes, followed by a 3X wash with PBS. Cells were then permeabilized with 0.1% Tween (KI67 imaging wells) or 0.3% Triton-X (total p53 imaging wells) in PBS with 1% BSA for 10min, followed by overnight incubation at room temperature with the primary antibody (ThermoFisher SolA15 for KI67, Cell Signaling Technology 7F5 for total p53). The following morning, cells were incubated with secondary antibody and imaged on an Opera Phenix.

### scRNA-seq and snRNA-seq QC, Filtering, and Annotation

Bcl2fastq was used to convert sequencing reads into bcl files for alignment with either the DropSeq pipeline (live-tissue scRNA-seq samples) or STARsolo (frozen-tissue snRNA-seq and HepG2 scRNA-seq samples). Mouse samples were aligned to the mm10 reference genome, and human samples were aligned to the Hg38 reference genome with the BFP sequence appended (to validate successful transduction via BFP positive control samples).

Cells were first filtered based on number of detected genes, detected UMIs, and percent of mitochondrial counts (Fig. S2-3, S6, S12, S15). The number of principal components was chosen based on an automated elbow-based selection criterion, followed by construction of nearest-neighbor and shared nearest-neighbor graphs and UMAP visualization. Clustering was implemented using the Leiden algorithm, and resolution was chosen based on a parameter scan and maximization of silhouette coefficient. Clusters solely distinguished by quality-associated or doublet-associated features were removed: 1) high expression of mitochondrial genes or well-established markers of low-quality cells (e.g., MALAT1, NEAT); or, 2) co-expression of markers for mutually-exclusive lineages (e.g., immune and epithelial cells). Cells were annotated based on canonical markers established in prior liver atlases, and clusters corresponding to broad lineages were merged for further subclustering (i.e., epithelial, immune, structural/stromal). Within each lineage, variable gene selection, PCA, nearest-neighbor graph construction, clustering, and UMAP visualization were re-performed; clusters distinguished by markers of other lineages were removed as doublets. Lineage-specific subclustering and doublet removal was repeated 2-3 times within each lineage to ensure robust heterotypic doublet identification.

### snATAC-seq QC, Filtering, and Annotation

CellRanger was used for processing and alignment to mm10-2020-A_arc_v2.0.0. Cells were filtered based on number of detected peaks, percent of reads in peaks, nucleosome signal, and TSS enrichment. The number of latent semantic indexing components was chosen based on an automated elbow-based selection, followed by construction of nearest-neighbor and shared nearest-neighbor graphs and UMAP visualization. Iterative subclustering and doublet removal were implemented as described in “scRNA-seq and snRNA-seq QC, Filtering, and Annotation”, using chromatin-based gene activity scores instead of canonical marker genes.

### Metabolic Adaptation Gene Program Derivation and Driver Gene Identification

At each timepoint for each of the scRNA-seq and snRNA-seq hepatocyte datasets, the average log_2_(fold-change) was calculated across all genes detected in at least 25% of cells (chosen based on differential expression benchmarking analyses examining robust parameter estimates and statistical outputs as a function of average expression and detection rate^192^). To promote prioritization of generalizable gene programs across species, we additionally incorporated differential expression information from Govaere et al.’s human bulk RNA-seq cohort, using limma-trend to model gene expression as a function of MASLD stage. See below for qualitative descriptions of the temporal patterns and trajectories captured by each gene program (defined quantitatively further below):

- Sustained Upregulation program: Genes whose elevation is maintained over time
- Sustained Downregulation program: Genes whose lessening is maintained over time
- Longitudinal Increase program: Genes progressively elevated with long-term chronic stress exposure
- Longitudinal Decrease program: Genes progressively lessened with long-term chronic stress exposure

Different genes are preferentially retained and measured in live-tissue scRNA-seq (nuclear + cytoplasmic mRNA) vs. frozen-tissue scRNA-seq (nuclear mRNA only). As a result, a gene may have a large fold-change in one dataset, but non-detection or sparse detection in the other dataset (in turn causing fold-changes that are inestimable or equal to 0). To maximize insights from both our live-tissue scRNA-seq dataset (higher molecular capture given the retention of cytoplasmic mRNA) and frozen-tissue snRNA-seq dataset (higher hepatocyte abundance), we sought to leverage complementary information from each dataset during gene program derivation. We followed the principle that genes should follow a given temporal trajectory (e.g., Sustained Upregulation) in at least one dataset, while allowing for non-detection/sparse detection in the other dataset (e.g., positive fold-changes in one dataset and non-negative fold-changes in the other dataset). As a result, gene programs were derived by filtering to retain genes matching the general structure shown below:

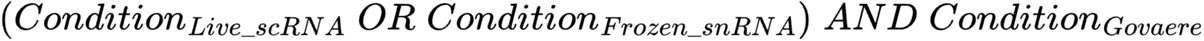

See below for each gene program’s definitions of Condition_Live_scRNA_, Condition_Frozen_snRNA_, and Condition_Govaere_, where log_2_FC represents average log_2_(fold-change) at the noted timepoint between hepatocytes from high fat vs. control diet mice in live-tissue scRNA-seq or frozen-tissue snRNA-seq datasets, and β_Govaere_limmatrend_ represents the regression coefficient of gene expression as a function of disease stage in Govaere et al.:

- Sustained Upregulation:

Condition_Live_scRNA_: Across all timepoints, consistent upregulation in live-tissue scRNA-seq and non-negative fold-changes in frozen-tissue snRNA-seq

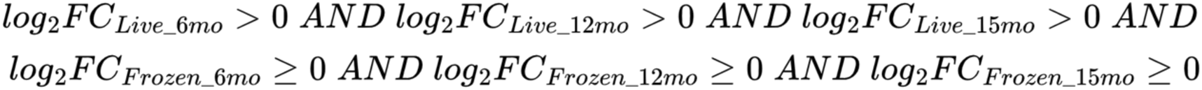
Condition_Frozen_snRNA_: Across all timepoints, consistent upregulation in frozen-tissue snRNA-seq and non-negative fold-changes in live-tissue scRNA-seq

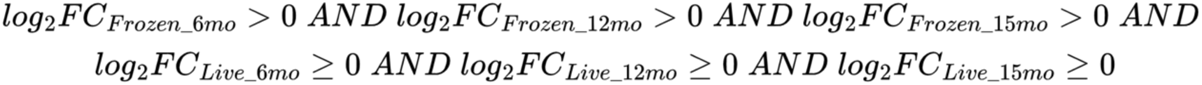
Condition_Govaere_: Increased expression with disease stage

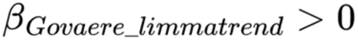
- Sustained Downregulation program:

Condition_Live_scRNA_: Across all timepoints, consistent downregulation in live-tissue scRNA-seq and non-positive fold-changes in frozen-tissue snRNA-seq

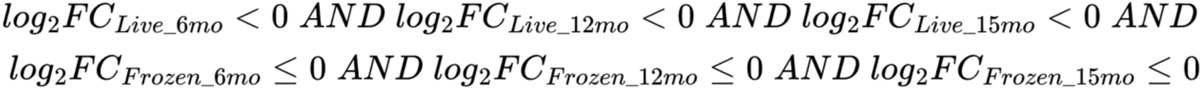
Condition_Frozen_snRNA_: Across all timepoints, consistent downregulation in frozen-tissue snRNA-seq and non-positive fold-changes in live-tissue scRNA-seq

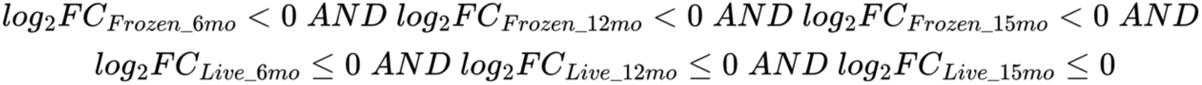
Condition_Govaere_: Decreased expression with disease stage

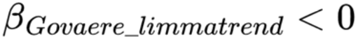
- Longitudinal Increase program:

Condition_Live_scRNA_: Progressive increases from 6-month to 15-month timepoints in live-tissue scRNA-seq and non-decreases in frozen-tissue snRNA-seq

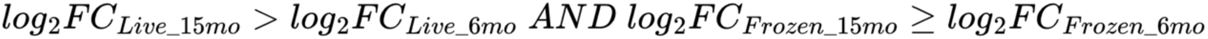
Condition_Frozen_snRNA_: Progressive increases from 6-month to 15-month timepoints in frozen-tissue snRNA-seq and non-decreases in live-tissue scRNA-seq

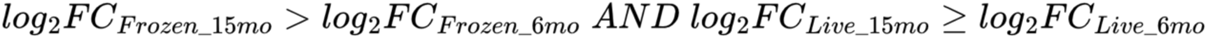
Condition_Govaere_: Increased expression with disease stage

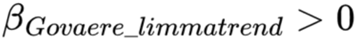
- Longitudinal Decrease program:

Condition_Live_scRNA_: Progressive decreases from 6-month to 15-month timepoints in live-tissue scRNA-seq and non-increases in frozen-tissue snRNA-seq

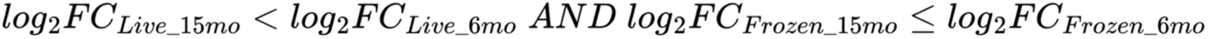
Condition_Frozen_snRNA_: Progressive decreases from 6-month to 15-month timepoints in frozen-tissue snRNA-seq and non-increases in live-tissue scRNA-seq

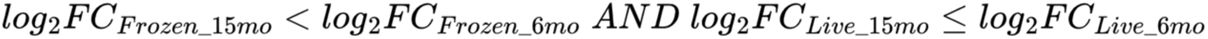
Condition_Govaere_: Decreased expression with disease stage

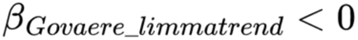

We highlight that only the mouse live-tissue scRNA-seq dataset, mouse frozen-tissue snRNA-seq dataset, and Govaere et al. bulk RNA-seq dataset were used for stress adaptation gene program derivation. Importantly, none of the other datasets presented elsewhere in the paper (e.g., Fig. 2P-Q, 3D-M, 6B-H, 7G-J, S3, S6, S10A, S7B-Q, or S12-14) were used as part of gene program derivation, ensuring that these figures’ analyses were conducted on “test sets” of entirely external, independent datasets. Module scores for each program and dataset were calculated using AddModuleScore in Seurat, and TCGA patient stratification and visualization were conducted with the survival and survminer packages in R.

### Comparison and Identification of Cancer Signature Induction during Stress Adaptation

To evaluate whether and which cancer and developmental phenotypes are mirrored in the spontaneous HCC tumors in our chronic metabolic stress mouse model, we began by conducting differential gene expression testing between tumor cells and hepatocytes in matched adjacent normal tissue. As a broad view of potential cancer phenotypes, we conducted gene set enrichment analysis using the fgsea R package, comparing our model’s tumor differential expression against gene sets from: 1) MSigDB’s “Chemical and Genetic Perturbations” (3,405 genesets spanning diverse prior studies’ mutational and signaling perturbations); 2) mutation-specific mouse liver tumor models^91^; and, 3) liver development and regenerative states^92–96^. Thus, all gene sets considered during gene set enrichment analysis were defined externally to this study, providing a complementary framework (to the previously-described adaptation program derivation) for uncovering phenotypes accentuated with tumorigenesis in our model.

We then sought to understand which tumorigenesis-linked gene sets were also differentially regulated during hepatocytes’ progressive stress adaptations (i.e., before tumorigenesis). Seurat’s AddModuleScore was used to score each dataset for leading-edge genes from statistically-significantly enriched cancer and developmental gene sets. Thus, statistically-significant module score differences in mouse tumor-vs-adjacent normal comparisons (Fig. 4A, 4F) reflect the statistically-significant gene set enrichment results on which the gene program is based; all significant differences in other datasets (Fig. 4B-E, 4G-J, S7D-E) represent “test set” results in entirely external datasets and independent extensions and connections to hepatocytes’ progressive adaptations to chronic stress across species and cohorts.

### Intercellular Signaling Analyses

We inferred potential intercellular signaling proteins that may drive hepatocytes’ stress adaptation gene programs by applying NicheNet to our live-tissue scRNA-seq data (which enables elevated proportions of immune cells as compared to frozen-tissue nuclei isolation-based datasets). Ligands were considered if detected in at least 10% of cells across any cell type. For each gene program, potential regulatory ligands were prioritized using NicheNet’s predict_ligand_activities. To control for ligands that broadly regulate hepatocyte functions or non-specific computational inference, we derived 100 random gene sets with matched average expression levels to the input gene program of interest; ligands’ regulatory potential scores were z-scored against a null background of their regulatory potentials for these random gene sets. Ligands were ranked by these regulatory potential z-scores, and only considered if their z-score was positive; a maximum of 8 ligands were retained for each gene program.

### Computational Methods for Longitudinal Epigenetic Alterations and Priming

Differential chromatin accessibility was calculated by pseudobulking snATAC-seq hepatocytes from each mouse, then calculating differential expression within each timepoint between high fat vs. control diet pseudobulked hepatocyte profiles using edgeR. chromVAR scores were calculated using the JASPAR2020 core motif collection and the Signac wrapper function RunChromVAR. Chromatin peak co-accessibility links were calculated using the Signac wrapper function run_cicero.

For higher-resolution inference of hepatocytes’ epigenetic adaptation trajectories, we conducted pseudotime analyses. To avoid confounding effects of diet and time, we derived separate pseudotime trajectories for hepatocytes from high fat and control diet hepatocytes, following concepts in previous analyses on allergic inflammation and stem cell differentiation^150^. Pseudotime trajectories were calculated for each diet condition using chromatin peaks that were differentially accessible with age within each diet condition. To control for varying cell counts across samples, each timepoint was downsampled to equal cell counts (matching the timepoint with the fewest cells). Latent semantic indexing and UMAP visualization (on LSI components 2 to 30) were implemented for each diet condition, and Slingshot was used to find pseudotime trajectories.

For epigenetic priming analyses, we began by linking distal peaks to genes based on co-accessibility with peaks in a gene’s promoter or gene body (only considering peaks within 100 kilobases of the gene and with Cicero co-accessibility score greater than 0.1). To identify genes whose putative distal chromatin regulators collectively exhibit differential accessibility, we then compared: 1) the observed distribution of differential accessibility fold-changes of actual co-accessible peaks; against, 2) 50 random background peaksets with matched average accessibility and GC bias. Finally, towards genes that may exhibit evidence of epigenetic priming in hepatocytes, we identified genes where:

1. Early chromatin accessibility changes exhibited the same directionality as transcriptional changes across longitudinal stress adaptation and tumorigenesis Epigenetic comparison: 6-month high fat vs. control diet snATAC-seq Transcriptional comparison: [15-month tumor vs. adjacent normal snRNA-seq] – [6-month high fat vs. control diet snRNA-seq]
2. Late chromatin accessibility changes (15-month high fat vs. control diet snATAC-seq) exhibited the same directionality as tumorigenesis transcriptional changes (15-month tumor vs. adjacent normal snRNA-seq) Epigenetic comparison: 15-month high fat vs. control diet snATAC-seq Transcriptional comparison: 15-month tumor vs. adjacent normal snRNA-seq
3. Tumorigenesis incurred a large shift in gene expression (15-month tumor vs. adjacent normal snRNA-seq) Transcriptional comparison: 15-month tumor vs. adjacent normal snRNA-seq
4. Early stress drove large changes in chromatin accessibility (6-month high fat vs. control diet snATAC-seq) Epigenetic comparison: 6-month high fat vs. control diet snATAC-seq differential accessibility fold-changes at gene-linked co-accessible peaks, relative to matched random background control

### MATCHA: Multiomic Ascertainment of Transcriptional Causality via Hierarchical Association

MATCHA seeks to prioritize transcription factors regulating arbitrary, user-specified gene programs, while leveraging multi-omic information on context-specific regulatory relationships (e.g., cell type- or tissue-specific gene-enhancer regulation).

As inputs, MATCHA accepts: 1) one or more arbitrary gene programs; 2) a TF motif database (e.g., JASPAR 2020); 3) multi-omic sc/snRNA-seq data; and optionally, 4) external datasets with relevance to the user’s context (e.g., prior bulk or sc/snRNA-seq atlases).

As outputs, MATCHA provides: 1) prioritization scores of the predicted strength and directionality of a TF’s regulatory effect on each arbitrary gene program (along with contributions of each input dataset to the overall prioritization score); 2) a bipartite network of which TFs regulate which gene programs (and in what direction, along with TF and gene program network centrality metrics); and, 3) rankings of which TFs may co-regulate multiple gene programs simultaneously (if multiple gene programs were provided).

Towards inference of gene program – co-accessible enhancer – causal TF triads, MATCHA is based on the principle that robust, strong regulatory relationships should be reflected across multiple -omic layers and across datasets. Therefore, MATCHA follows the following steps:

1. If a particular TF regulates the user-specified gene program(s), then it is plausible that the regulatory relationship should be reflected in accessibility changes at program-associated chromatin regions containing the TF’s motif. Therefore, MATCHA begins by identifying chromatin regions that are co-accessible with a gene’s promoter or gene body (based on Cicero), then filters peaks based on genomic distance and co-accessibility strength towards plausible regulatory relationships. For each TF motif, MATCHA creates a program-specific motif score by evaluating accessibility at program-coaccessible peaks containing each TF motif (based on chromVAR). Finally, MATCHA calculates the correlation between the program-specific motif score and transcriptional expression of the gene program itself. In this way, MATCHA connects epigenetic alterations at program-linked, TF motif-containing peaks to transcriptional levels of the gene program.
2. If a particular TF regulates the user-specified gene program(s), then it is plausible that the regulatory relationship should be reflected in concordant changes between TF abundance and gene program transcriptions. Therefore, MATCHA calculates the correlation between expression level of each TF in the user-input TF motif database and transcriptional expression of the gene program, thereby connecting transcriptional TF abundance to transcriptional levels of the gene program.
3. If a particular TF regulates the user-specified gene program(s) robustly, then it is plausible that the regulatory relationship should be reflected not only in a single dataset, but across studies. Therefore, if the user provided multiple input datasets (whether bulk or single-cell), MATCHA calculates similar correlations as described above between gene program transcriptional level and either TF motif accessibility at program-linked peaks or TF transcriptional abundance. In this way, MATCHA connects each TF (motif) to transcriptional levels of the gene program across wide-ranging contexts (e.g., spanning species, disease severities, experimental designs, measurement technologies, etc.).
4. To create a single prioritization score for each TF that incorporates information across - omic measurements and studies, MATCHA aggregates each correlation (as calculated in preceding steps). To account for different correlation distributions across -omic measurement modalities and studies, correlations calculated in each previous step are scaled from −1 (most negative association between TF and gene program, indicative of repression) to 1 (most positive association between TF and gene program, indicative of activation). Scaled correlations are averaged, and TFs are ranked by the average of scaled correlations.
5. In addition to identifying which TFs regulate a particular gene program, it is important to understand whether TFs may co-regulate multiple gene programs, towards essential couplings of cellular phenotypes encapsulated by different programs or potential opportunities to disentangle otherwise-associated phenotypes. Therefore, if the user provided multiple input gene programs, MATCHA will create rankings of each TF’s cross-dataset association with each gene program (as described in preceding steps), and calculate network centrality metrics for TFs and gene programs (e.g., out-degree for TFs to identify the number of gene programs that they strongly regulate, in-degree for gene programs to identify the number of TFs potentially regulating them). For concise summarization, MATCHA will filter to retain TFs that are ranked within the top TFs in at least a minimum number of gene programs (Fig. 5F generated with the top 10 TFs for each gene program, showing TFs linked to at least 2 gene programs). Optionally, users can further filter to retain only TFs whose regulatory relationships match observed transcriptional correlations between gene programs (e.g., if a given TF is linked to program_1_ and program_2_ which are in turn negatively correlated with each other at the transcriptional level, the TF must activate one of the programs but repress the other). In this way, MATCHA enables nomination and prioritization of TFs with strong regulatory effects on wide-ranging, but specific cellular phenotypes of particular interest to the user (e.g., this work’s stress adaptation gene programs, with distinct temporal trends, functional enrichments, and prognostic stratification of human HCC survival).

## Acknowledgements

We thank Maria Alimova in the Center for the Development of Therapeutics, for her expertise with high-throughput imaging. We thank the Swanson Biotechnology Center at the Koch Institute, which encompasses the Flow Cytometry, Histology, and Genomics & Bioinformatics Core facilities (NCI P30-CA14051). We thank the Department of Comparative Medicine for mouse husbandry support. We thank Sven Holder and members of the Hope Babette Tang (1983) Histology Facility for histology support. C.N.T. is supported by a Fannie and John Hertz Foundation fellowship and a National Science Foundation Graduate Research Fellowship (1122374). M.S.S is supported by the National Institutes of Health Grant T32DK007191. J.E.S. is supported by a National Institutes of Health F32 Fellowship (F32DK128872). Ö.H.Y. is supported by National Institutes of Health Grant R01CA245314. W.G. is supported by National Institutes of Health Grants R01DK090311, R01DK105198, R24OD017870. C.N.T., J.E.S, Ö.H.Y., and A.K.S receive support from the MIT Stem Cell Initiative through Fondation MIT.

## Data and materials availability

All code used for scRNA-seq analysis are accessible at Zenodo under XX. MATCHA is available on Github for download and use at XX. scRNA-seq digital gene expression matrices, metadata, and interactive visualization tools can be found as study XX through the Alexandria Project, a Bill & Melinda Gates Foundation–funded portal (part of the Single Cell Portal hosted by the Broad Institute of MIT and Harvard). FASTQs have been uploaded to GEO at accession number XX.

