## Supplemental Information for "Chronic metabolic stress drives developmental programs and loss of tissue functions in non-transformed liver that mirror tumor states and stratify survival"

### Supplementary Note 1: Contextualization of diet-only chronic stress mouse model against human metabolic stress adaptation progression

We additionally sought to contextualize phenotypes observed in the diet-only mouse model used in this work to those observed in human patients with the progression of MASLD and HCC. H&E and trichrome stains were blindly scored by a clinical liver pathologist[1], revealing that HFD mice typically demonstrate simple steatosis at 6 months and MASLD by 12 months (Fig. S1). HFD livers developed classic ballooning degeneration by 12 months, often associated with human patients' progression to more severe MASLD (Fig. 1G-H)[2]. Hepatocyte ballooning was confirmed by a second clinical liver pathologist and further validated by negative staining of CK8/18[3,4] (Fig. 1H). These temporal patterns of steatosis and fibrosis align with the well-known "burnt-out NASH" and "cryptogenic cirrhosis" phenotypes where hepatic fat is lost in human patients with end-stage liver disease[5–7].

Supplementary Note 2: Connections between hepatocytes' stress adaptation programs, acute regeneration phenotypes, and intercellular signaling drivers

Towards connections to the liver's intrinsic regenerative capacity, we found that hepatocytes undergoing acute regeneration exhibit temporally-coordinated, internally-consistent expression patterns of the Longitudinal Increase, Longitudinal Decrease, Development-Associated, and Human HCC S1: WNT Activation programs derived or highlighted in this work, in line with hepatocytes' adaptations to metabolic stress drawing upon the liver's regenerative capabilities (Fig. S9J)[8].

To more holistically examine intercellular signaling that may contribute to hepatocyte phenotypic alterations, we leveraged our immune-biased live tissue scRNA-seq dataset to infer ligands poised to regulate hepatocytes' metabolic adaptations (based on NicheNet; see Methods). Among the ligands predicted to regulate stress adaptation programs associated with increased disease severity and worsened tumor survival were neutrophil-derived MMP9, Kupffer cell-derived GDF3 and RTN4, endothelial cell-derived CTGF and EDN1, and T cell-derived LTB (Fig. S9F-I). All of these ligands exhibited upregulated expression or increased sender cell abundances with high fat diet, complementing their literature associations with worsened MASLD severity, increased regeneration, and/or tumorigenesis contributions[9–15]. Intercellular signaling interactions nominated through our analyses, but with comparatively less literature support, may represent promising directions for future experimental validation of cell-extrinsic mediators of hepatocyte dysfunction.

Supplementary Note 3: Contextualization of computational methods to infer transcription factors driving arbitrary gene programs

While a wide variety of computational methods exist to estimate TF abundance or activity, the goal of identifying TFs that specifically regulate particular phenotypic gene programs remains an open problem. Below, we highlight generally-related approaches and distinctions from MATCHA's goal of mapping gene programs, enhancers, and program-specific TFs. We also note that we do not intend the following descriptions to represent a comprehensive or all-encompassing review (e.g., see [16–19] for examples of prior reviews and benchmarking efforts).

- Examining differentially-expressed TFs (at the transcriptional level) or genome-wide TF motifs with differential accessibility (at the chromatin level) can prioritize TFs with generally altered activity, but not connect them to specific cellular phenotypes.
- Genome-wide transcriptomic profiles can be translated into predicted TF activities through tools that leverage prior knowledge (e.g., response signatures, inferred interactomes, etc.; see [20–23] for examples). However, these tools provide a single TF score for each transcriptomic profile, but do not associate TFs with specific programs or distinguish their effects (or lack thereof) on specific cellular phenotypes.
- Construction of gene regulatory networks can connect transcription factors to inferred downstream genes (with optional incorporation of multi-omic information)[17,24–27]. However, these tools most often focus on inference of TFs distinguishing distinct cell types (e.g., differentiation from one developmental stage to another), or simulated transitions across genome-wide transcriptomic profiles (e.g., represented as regions of density on a low-dimensional visualization such as UMAP or t-SNE). Thus, frameworks to connect TF regulons to user-specified gene programs of interest remain less clear. While gene regulatory network inference connects TFs to potential targets, the ability to define gene programs of interest and prioritize causal TFs has received less exploration. It remains difficult to identify TFs that target and modulate specific disease-associated phenotypes, rather than affecting the global cell state.

In contrast, MATCHA begins from a user-specified gene program and enables prioritization and nomination of TFs regulating that particular gene program, towards TFs driving specific aspects of cellular physiology. In this way, MATCHA enables discovery of tradeoffs or co-regulation across disease-linked axes of variation with distinct functions and repercussions (e.g., this work's Longitudinal Increase and Decrease programs, with opposing enriched processes, temporal trajectories, and disease connections). We highlight that MATCHA: 1) leverages cell type- and/or tissue-specific transcriptomic and epigenetic relationships, towards capturing context-specific regulatory interactions, 2) can emphasize robust and generalizable gene program drivers by incorporating datasets spanning studies, cohorts and species, and, 3) can uncover core, co-regulatory TFs capable of activating or repressing each of multiple gene programs simultaneously.

Supplementary Note 4: Contextualization of p53 regulation of hepatocyte phenotypes

In the liver, p53 is known to have roles beyond its canonical anti-cancer genoprotective and antiproliferative functions. For instance, p53 modulates a variety of metabolic pathways (e.g., lipid processing, gluconeogenesis) in the liver, suggesting potential additional relevance for this TF in the context of metabolic overload[28,29]. Towards metabolic functions of p53 during lipid overload, liver knockout of p53 in mice can produce steatosis, and p53 upregulation (via degradation of an upstream inhibitor) can alternatively attenuate fat accumulation[30]. Additionally, elevated p53 activity has also been linked to progenitor-associated hepatocyte phenotypes. Multiple studies have shown that p53 can repress hepatocytes' lineage-determining transcription factor HNF4A, across reductions of HNF4A protein levels and promoter activity[31,32]. p53 stabilization (e.g., via MDM2 deletion) can induce progenitor markers in vivo; proposed mechanisms include p53 inducing inflammation that in turn promotes progenitor states or microRNA-mediated inhibition of HNF4A[33,34].

As literature support for KI67 accumulation and hepatocyte cycling despite elevated p53, both p53 and KI67 can increase with HCC grade. As specific examples, these genes were upregulated in tumors from a cohort of HBV-associated HCC patients relative to paired normal liver tissue[35]; in a separate human HCC cohort, p53 levels correlated with the abundance of PCNA (another common cell cycle marker), and each associated with worse tumor grade[36]. Additionally, in liver regeneration models using *in vitro* primary human hepatocytes and *in vivo* mouse livers, WNT signaling activation and AP-1 transcription factors can each suppress p53's anti-proliferative effects and abrogate p53-mediated cell cycle inhibition in hepatocytes[37,38]; our transcriptional and epigenetic datasets support increased activities of each of these pathways over the course of MASLD progression. Thus, in particular cases in the liver, p53 and KI67 are not necessarily anti-correlated and can exhibit concordant regulation upon cellular perturbations.

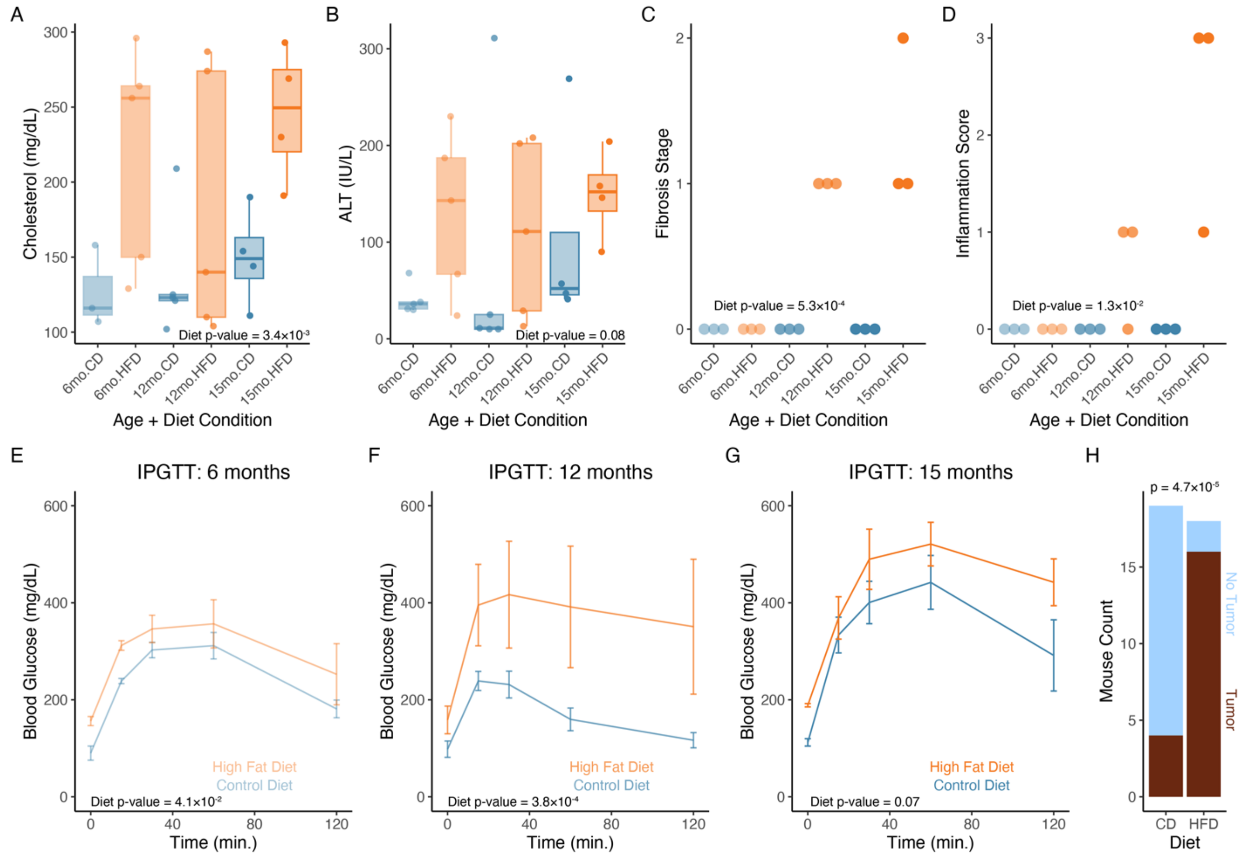


Supplementary Figure 1: Characterization of mouse functional, organ-level, and tissue-level phenotypes induced by metabolic stress. (A-D) Circulating alanine transaminase concentrations (A), circulating cholesterol (B), histologic fibrosis stage (C), and histologic inflammation score (D) across chronic metabolic stress progression; each dot is an independent mouse. (E-G) Intraperitoneal glucose tolerance test across diet conditions at 6-month (E), 12-month (F), and 15-month (G) timepoints (N = 3-5 per age×diet condition); bars indicate standard error of the mean. (H) Spontaneous liver tumor occurrence counts in orthogonal HFD mouse validation cohort at an independent mouse facility. P-values in (A-G) calculated through two-way ANOVA; p-value in (H) calculated through Fisher’s exact test.

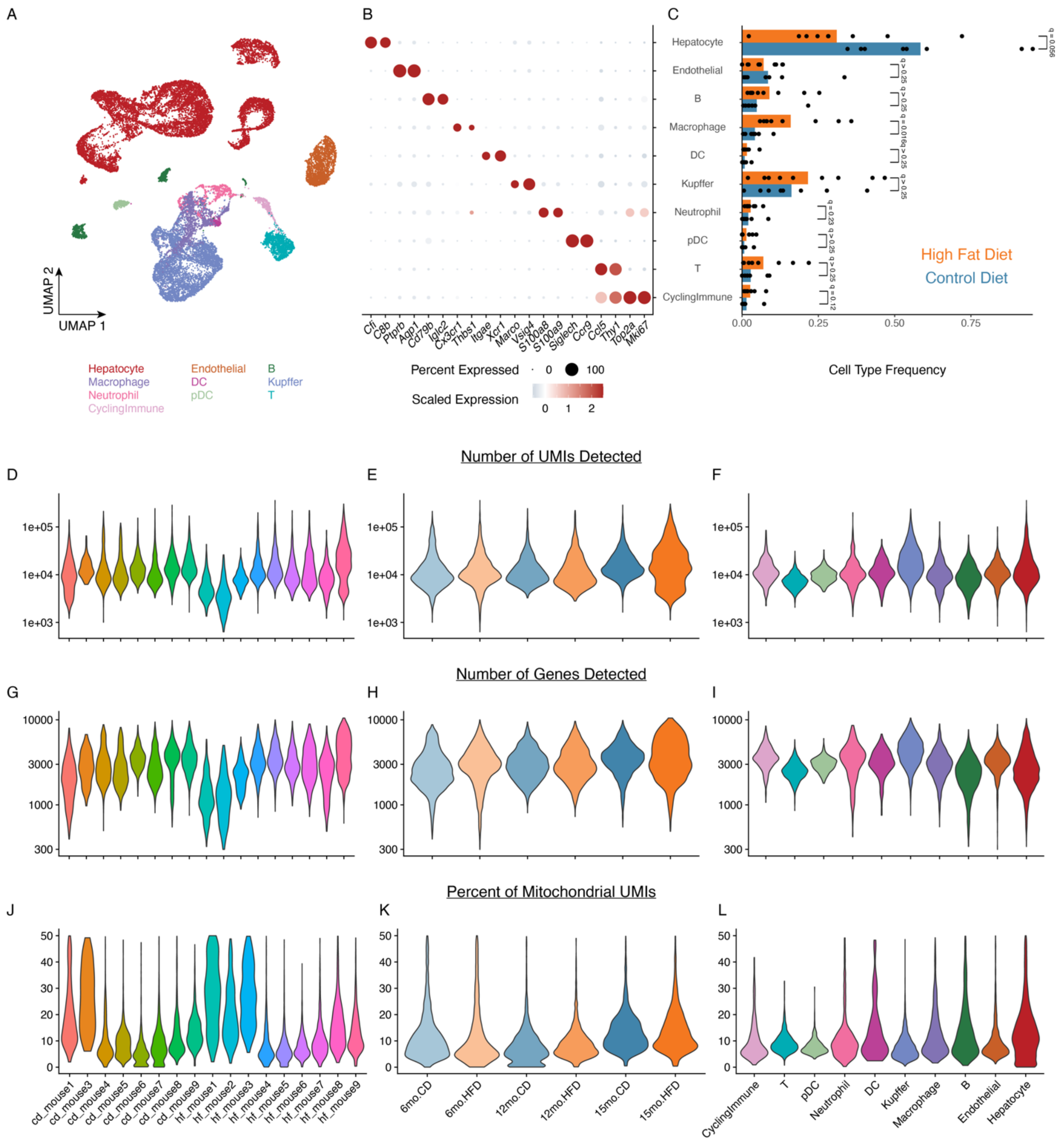


<u>Supplementary Figure 2: Data overview and QC metrics of live-tissue scRNA-seq.</u> (A) UMAP visualization of live-tissue scRNA-seq data. (B) Marker gene dotplot visualization and (C) compositional abundance of cell types. (D-F) Number of detected UMIs split by (D) mouse, (E) age and diet condition, or (F) cell type. (G-I) Number of detected genes split by (G) mouse, (H) age and diet condition, or (I) cell type. (J-L) Percent of mitochondrial UMIs split by (J) mouse, (K)

age and diet condition, or (L) cell type. P-values in (C) calculated through Mann-Whitney U-test with Benjamini-Hochberg multiple testing correction.

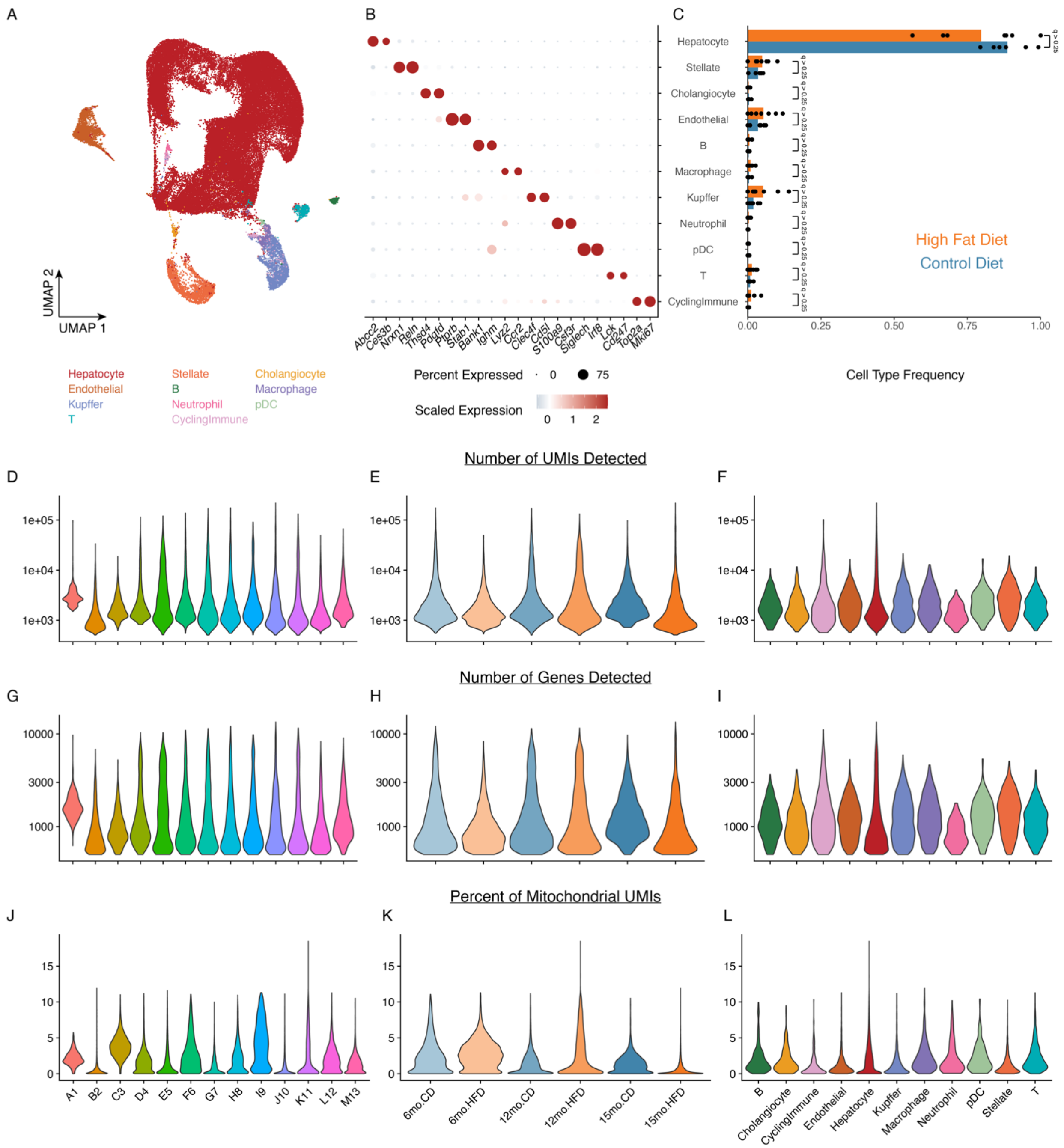


<u>Supplementary Figure 3: Data overview and QC metrics of frozen-tissue snRNA-seq.</u> (A) UMAP visualization of frozen-tissue snRNA-seq data. (B) Marker gene dotplot visualization and (C) compositional abundance of cell types. (D-F) Number of detected UMIs split by (D) mouse, (E) age and diet condition, or (F) cell type. (G-I) Number of detected genes split by (G) mouse, (H) age and diet condition, or (I) cell type. (J-L) Percent of mitochondrial UMIs split by (J) mouse, (K)

age and diet condition, or (L) cell type. P-values in (C) calculated through Mann-Whitney U-test with Benjamini-Hochberg multiple testing correction.

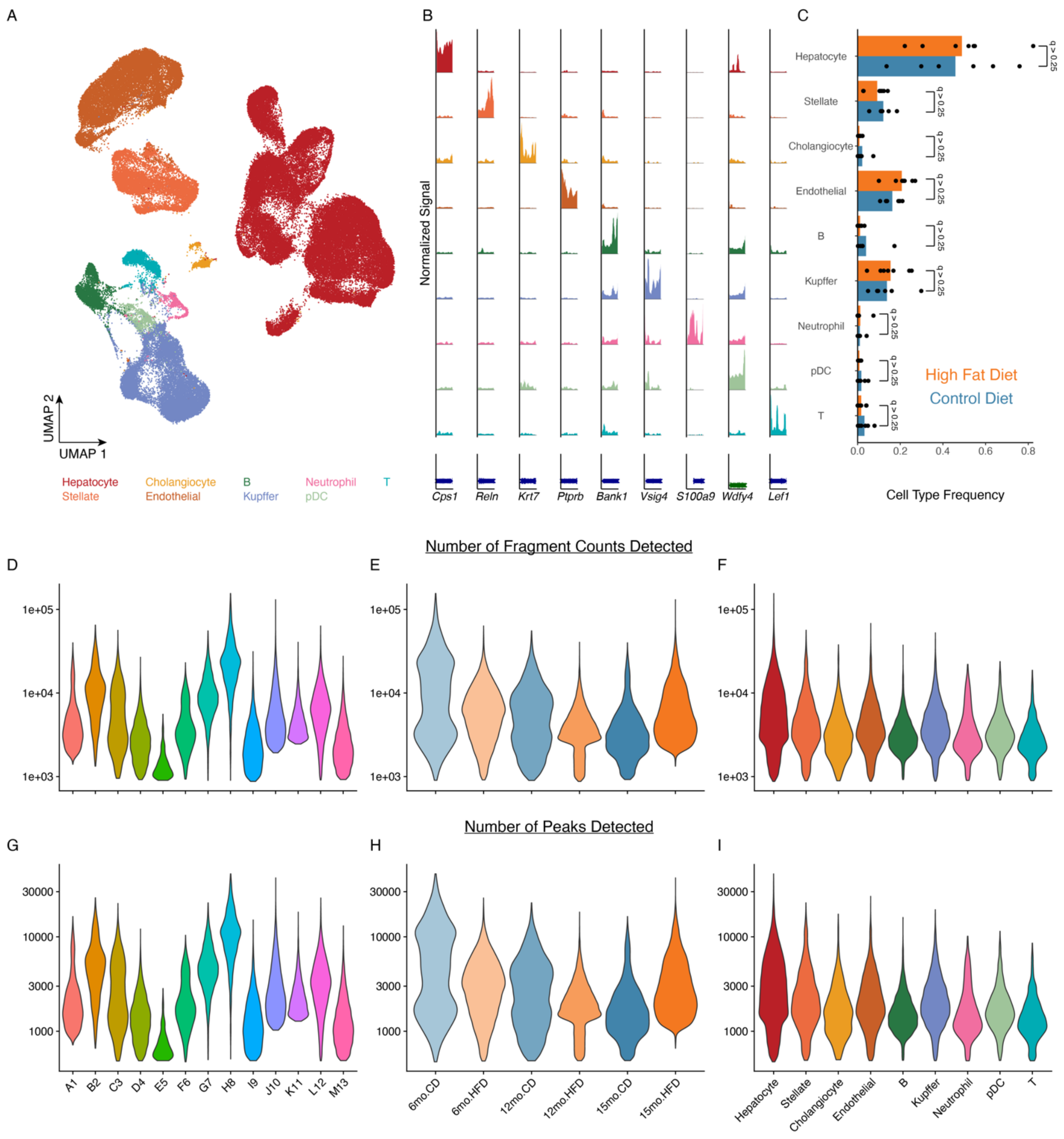


Supplementary Figure 4: Data overview and QC metrics of frozen-tissue snATAC-seq. (A) UMAP visualization of frozen-tissue snATAC-seq data. (B) Chromatin accessibility coverage plate visualization and (C) compositional abundance of cell types. (D-F) Number of detected fragment counts split by (D) mouse, (E) age and diet condition, or (F) cell type. (G-I) Number of detected peaks split by (G) mouse, (H) age and diet condition, or (I) cell type. P-values in (C) calculated through Mann-Whitney U-test with Benjamini-Hochberg multiple testing correction.

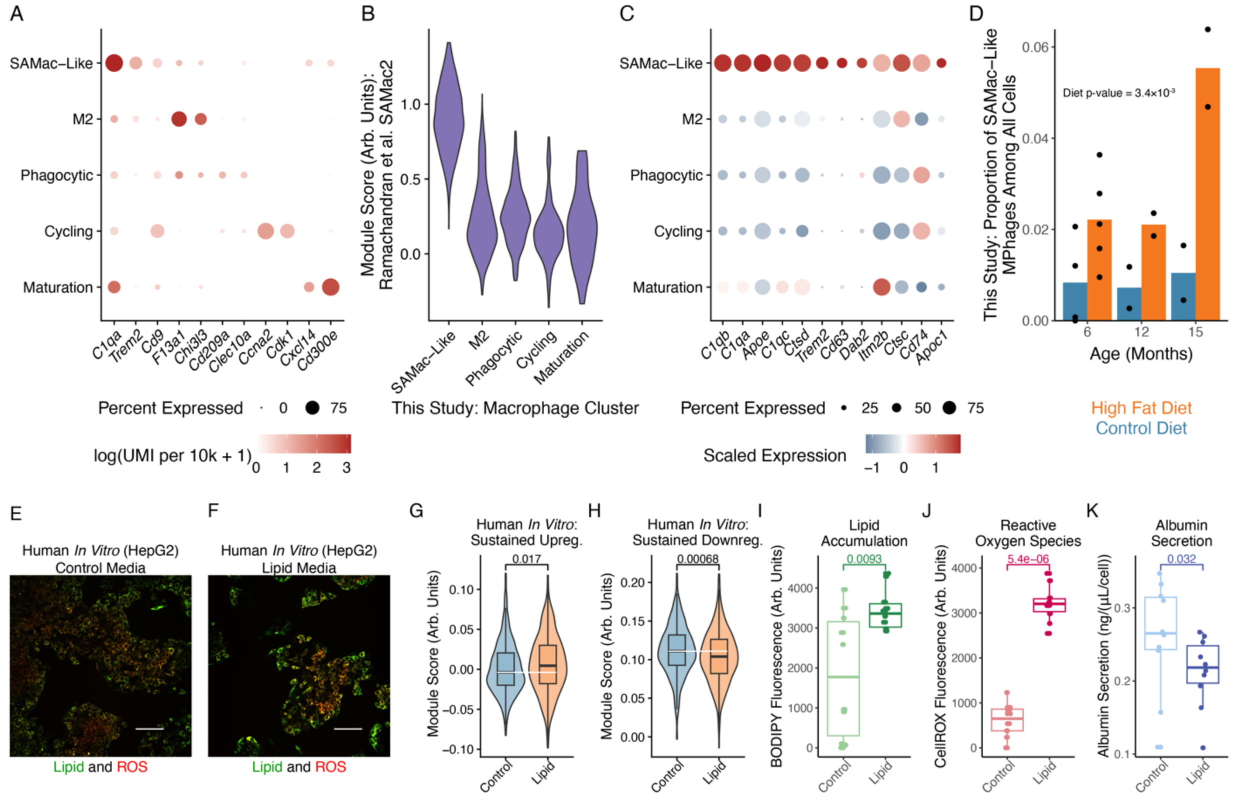


<u>Supplementary Figure 5: Mouse macrophage subcluster exhibits transcriptional similarities to human SAMac cells.</u> (A) Dotplot of marker genes distinguishing this study's macrophage subclusters. (B) Module score of SAMac2 markers, split by macrophage subcluster. (C) Dotplot of SAMac marker genes highly expressed in this study's SAMac-like subcluster. (D) Compositional enrichment of SAMac-like macrophages with high fat diet even at initial timepoint. (E-F) HepG2 lipid accumulation (BODIPY 493) and ROS accumulation (CellROX) in control media (E) or lipid media (F) (scalebar=100µm). (G-H) Sustained Upregulation program (G) and Sustained Downregulation program (H) expression in HepG2 cells (n = 891 cells). (I-K) HepG2 lipid accumulation (I; microscopy), ROS accumulation (J; microscopy), albumin secretion (K; ELISA). Dots indicate independent wells; N = 10 wells/condition. P-value in (D) calculated through two-way ANOVA; all other p-values calculated through Mann-Whitney U-test.

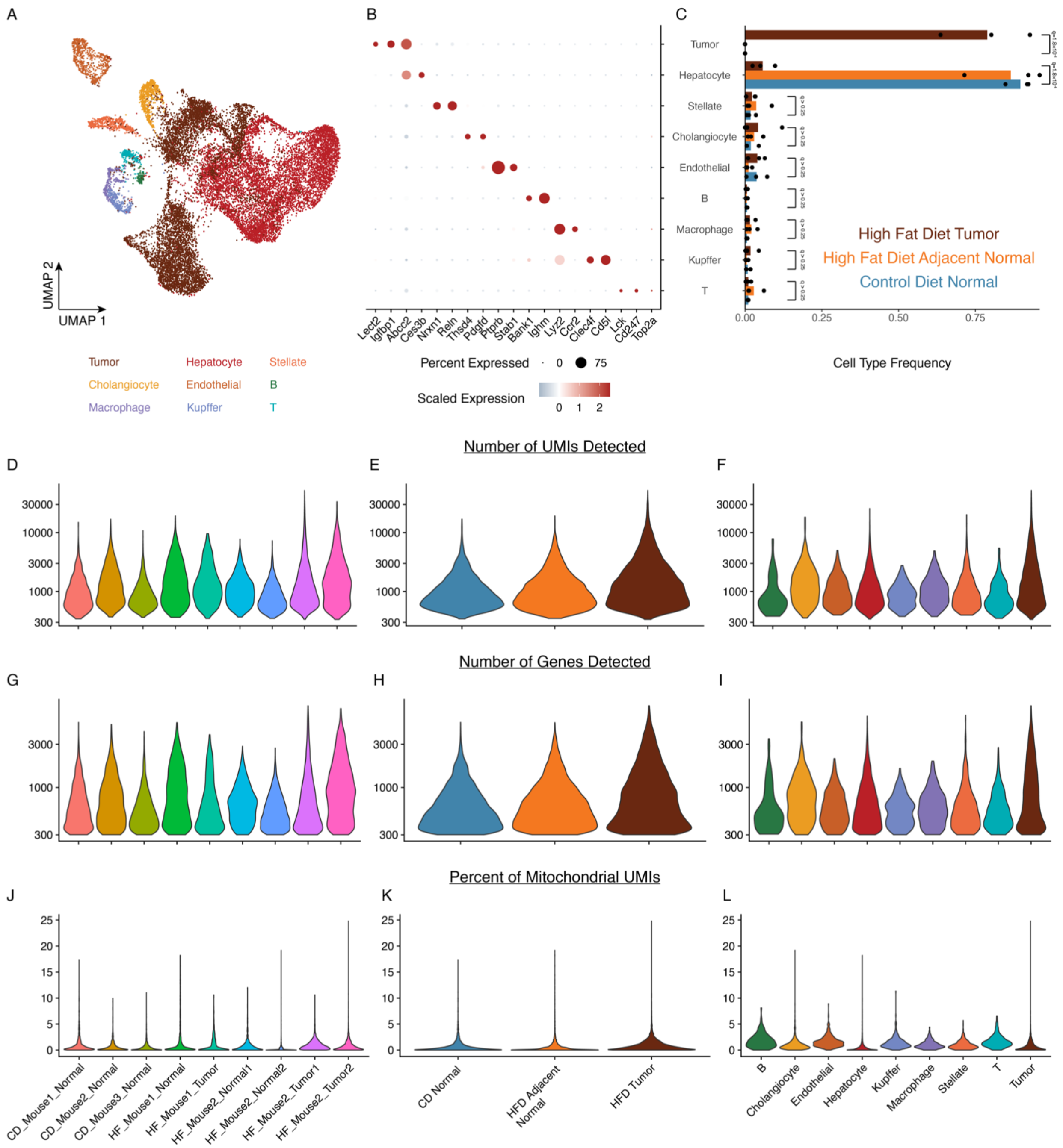


<u>Supplementary Figure 6: Data overview and QC metrics of frozen-tissue tumor snRNA-seq.</u> (A) UMAP visualization of frozen-tissue tumor snRNA-seq data. (B) Marker gene dotplot visualization and (C) compositional abundance of cell types. (D-F) Number of detected UMIs split by (D) mouse, (E) diet and tumor status, or (F) cell type. (G-I) Number of detected genes split by (G) mouse, (H) diet and tumor status, or (I) cell type. (J-L) Percent of mitochondrial UMIs split by (J) mouse, (K) diet and tumor status, or (L) cell type. P-values in (C) calculated through one-way ANOVA.

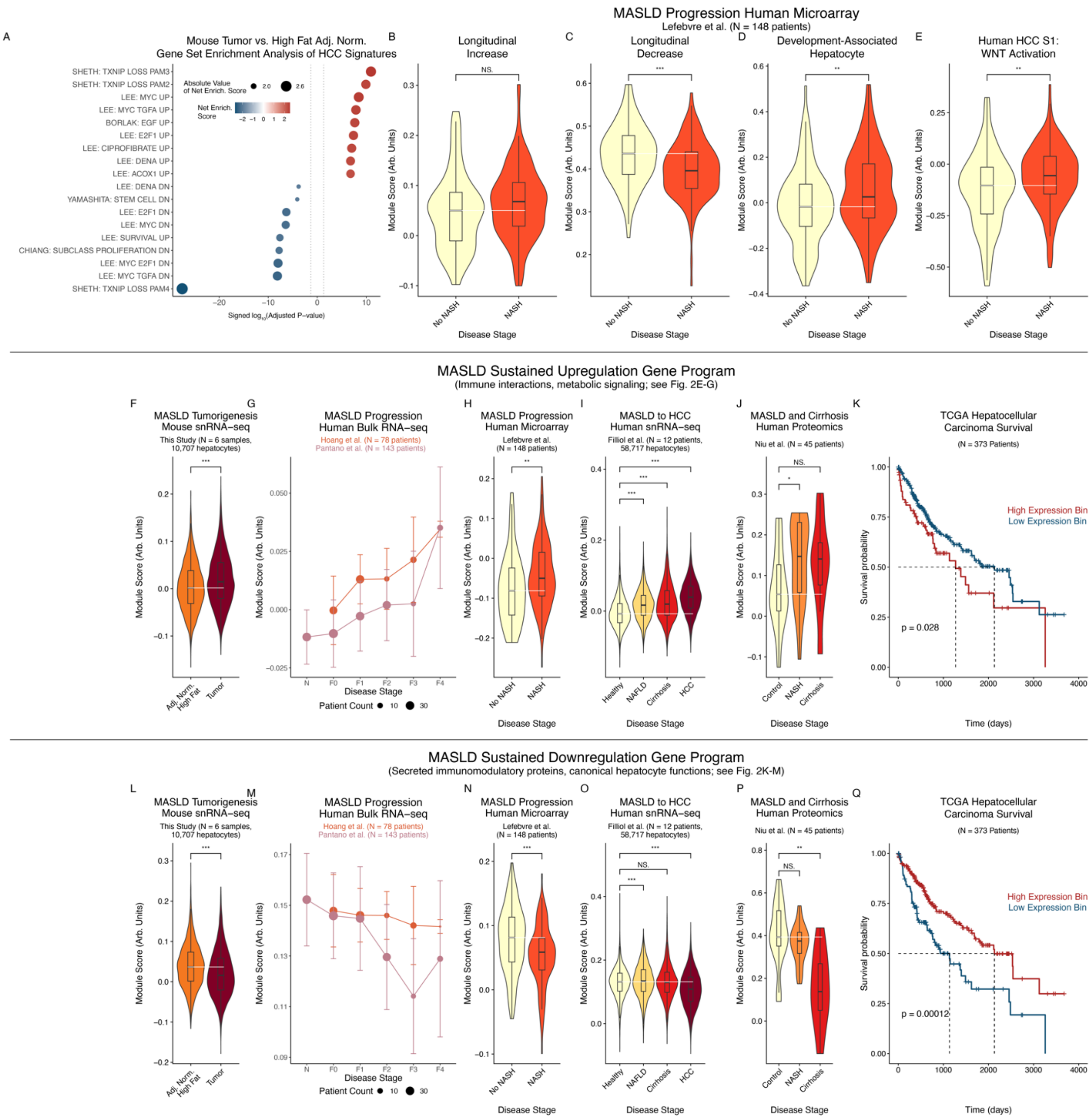


<u>Supplementary Figure 7: Metabolic stress adaptation programs across species, cohorts, measurements, and clinically-relevant outcomes.</u> (A) Gene set enrichment analysis of HCC signatures among differentially-expressed genes between mouse high fat diet tumor and adjacent normal samples. (B-E) Expression in external human MASLD cohort (liver bulk microarray) of (B) Longitudinal Increase program, (C) Longitudinal Decrease program, (D) development-associated hepatocyte program, or (E) Human HCC S1: WNT Activation program. (F) Expression of Sustained Upregulation program in tumor cells and hepatocytes from adjacent

normal tissue (15-month HFD). (G) Expression of Sustained Upregulation program in bulk RNA-seq from external human MASLD cohorts. (H) Expression of Sustained Upregulation program in liver microarray from external human MASLD cohort. (I) Expression of Sustained Upregulation program in hepatocytes and tumor cells from snRNA-seq from external human MASLD/HCC cohorts. (J) Protein abundance of Sustained Upregulation program in proteomics from external human MASLD/cirrhosis cohort. (K) Stratification of TCGA HCC patient survival, based on expression level of Sustained Upregulation program. (L-Q) Disease progression, cancer phenotype associations, and predictive power of Sustained Downregulation program, following format of (F-K). Survival outcome p-values calculated with log-rank test; GSEA net enrichment score and p-value calculated using random permutation testing through fgsea package; all other p-values calculated using Mann-Whitney U test.

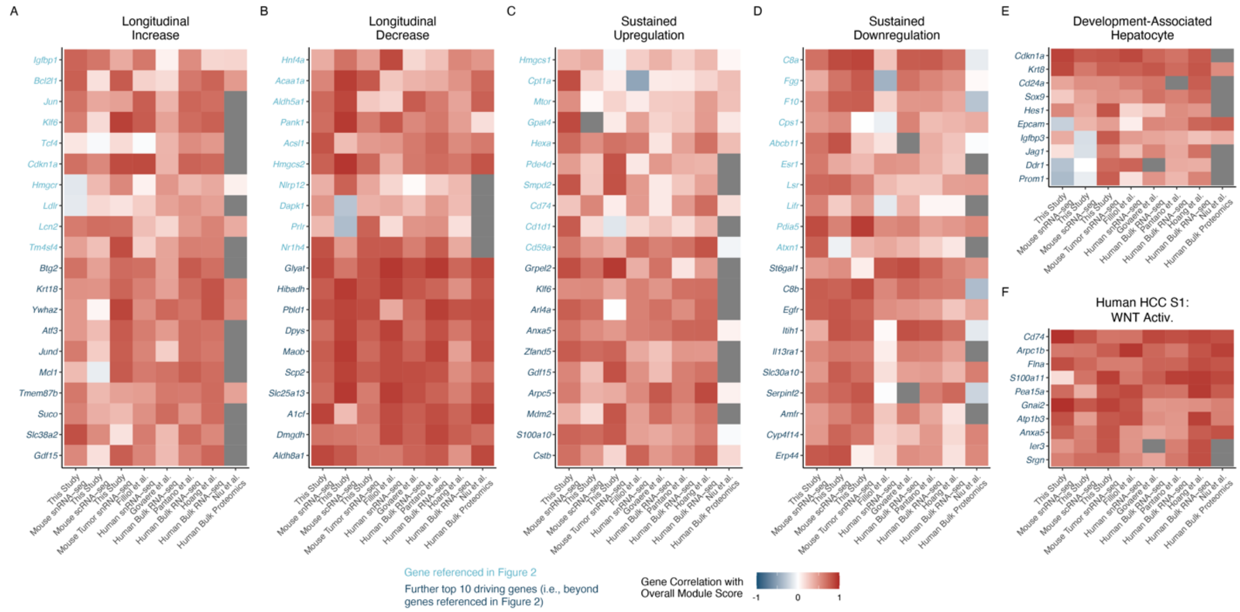


Supplementary Figure 8: Driver genes of overall program scores. Spearman correlation of individual genes with consistent, strong, positive covariance with overall program scores across datasets: (A) Longitudinal Increase program, (B) Longitudinal Decrease program, (C) Sustained Upregulation program, (D) Sustained Downregulation program, (E) Development-Associated Hepatocyte program, (F) Human HCC S1: WNT Activation program.

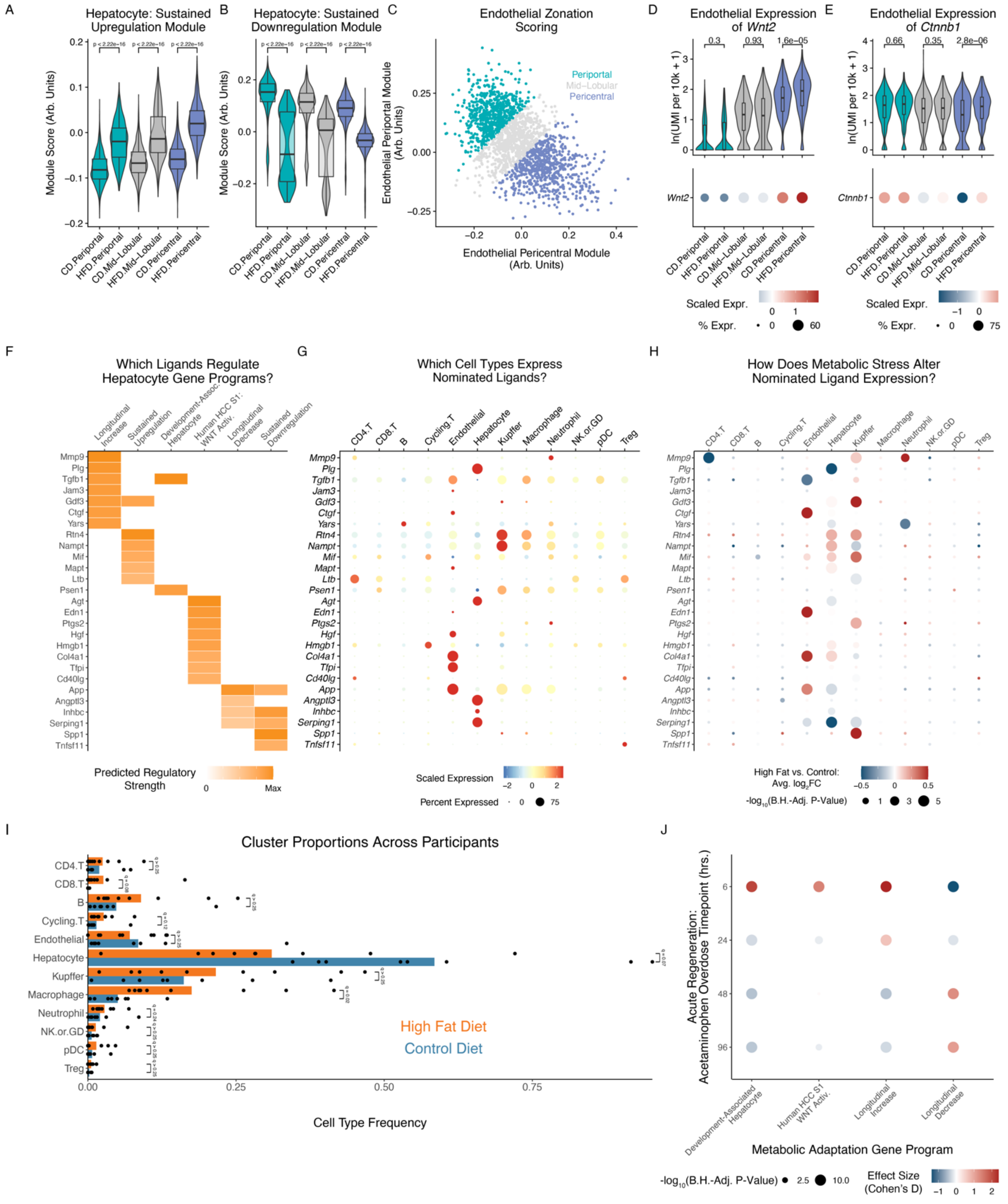


<u>Supplementary Figure 9: Connection of stress adaptation programs to intercellular signaling and acute regeneration.</u> (A-B) Aggregate hepatocyte module score of (A) Sustained Upregulation or (B) Sustained Downregulation program expression levels, over diet and inferred spatial zonation.

(C) Zonation scoring and binning of endothelial cells from our mouse model. (D-E) Expression of (D) *Wnt2* or (E) *Ctnnb1* in endothelial cells, over diet and inferred spatial zonation. (F) Top inferred ligands prioritized as regulating hepatocytes' stress adaptation-associated gene programs. (G) Average expression level of inferred ligands across cell types in our live tissue scRNA-seq mouse data. (H) Change in ligand expression with metabolic stress across cell types in our live tissue scRNA-seq mouse data. (I) Change in cell type abundances with metabolic stress in our live tissue scRNA-seq mouse data. (J) Effect size of change in expression levels of this work's stress adaptation programs during acute regeneration (comparing timepoints after acetaminophen overdose to expression at t = 0). All p-values calculated using Mann-Whitney U test.

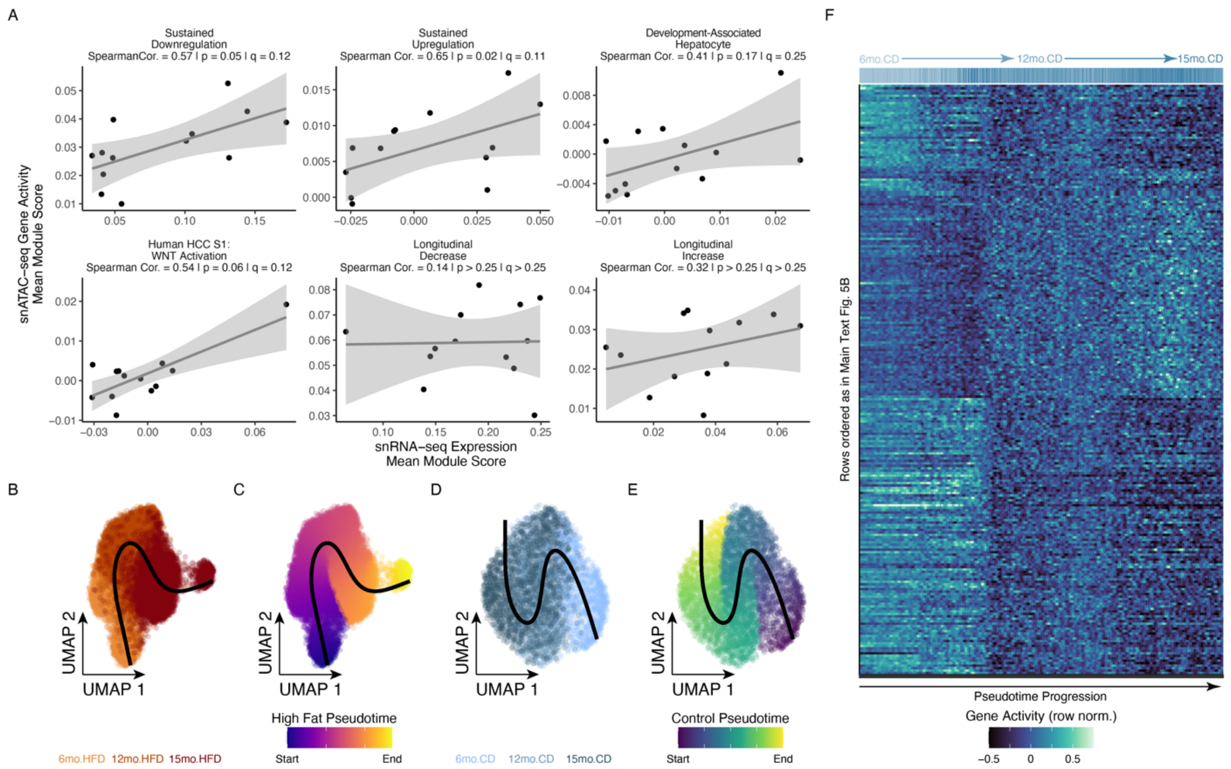


Supplementary Figure 10: Epigenetic trajectories of hepatocytes adapting to chronic metabolic stress. (A) Association between transcriptomic expression (x-axis) and chromatin accessibility (y-axis) of gene programs uncovered in this work (pseudobulked tandem snRNA-seq and snATAC-seq datasets). (B-C) Low-dimensional embedding and visualization of high-fat diet snATAC-seq hepatocytes, colored by (B) timepoint or (C) position along inferred pseudotime progression. (D-E) Low-dimensional embedding and visualization of control diet snATAC-seq hepatocytes, colored by (D) timepoint or (E) position along inferred pseudotime progression. (F) Heatmap of chromatin accessibility-based gene activity scores in control diet hepatocyte pseudotime progression; rows have matched order with Main Text Fig. 5B.

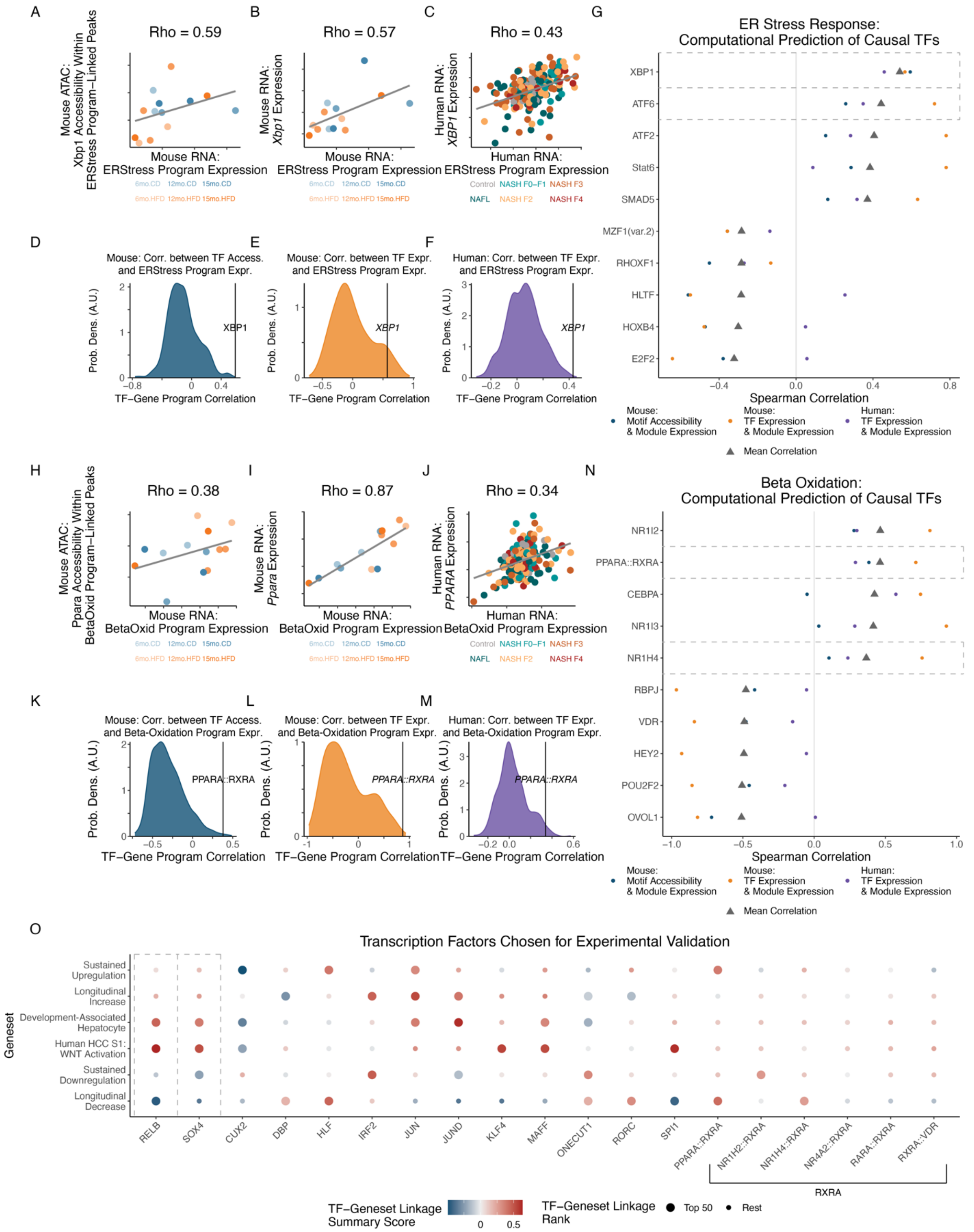


Supplementary Figure 11: MATCHA enables prioritization of TFs regulating arbitrary user-defined gene programs and specific cellular phenotypes. (A) Spearman correlation between transcriptomic expression of the ER stress response gene program (defined externally to this study through the public GO:BP database) and Xbp1 motif accessibility within program-coaccessible linked peaks (this study's natural progression tandem snRNA-seq and snATAC-seq datasets). (B) Spearman correlation between transcriptomic expression of the ER stress response gene program and Xbp1 transcriptomic expression (this study's natural progression snRNA-seq datasets). (C) Spearman correlation between transcriptomic expression of the ER stress response gene program and XBP1 transcriptomic expression (Govaere et al.'s human patient cohort). (D) Distribution of Spearman correlations between ER stress response gene program transcriptomic expression and accessibility of all TF motifs at program-coaccessible linked peaks (this study). (E) Distribution of Spearman correlations between ER stress response gene program transcriptomic expression and expression of all TFs (this study). (F) Distribution of Spearman correlations between ER stress response gene program transcriptomic expression and expression of all TFs (Govaere et al.). (G) Maximally prioritized TFs inferred as driving (top 5) or repressing (bottom 5) ER stress response gene program. (H) Spearman correlation between transcriptomic expression of the beta-oxidation response gene program (defined externally to this study through the public GO:BP database) and PPARA::RXRA motif accessibility within program-coaccessible linked peaks (this study's natural progression tandem snRNA-seq and snATAC-seq datasets). (I) Spearman correlation between transcriptomic expression of the beta-oxidation gene program and Ppara transcriptomic expression (this study's natural progression snRNA-seq datasets). (J) Spearman correlation between transcriptomic expression of the beta-oxidation gene program and PPARA transcriptomic expression (Govaere et al.'s human patient cohort). (K) Distribution of Spearman correlations between beta-oxidation gene program transcriptomic expression and accessibility of all TF motifs at program-coaccessible linked peaks (this study). (L) Distribution of Spearman correlations between beta-oxidation gene program transcriptomic expression and expression of all TFs (this study). (M) Distribution of Spearman correlations between beta-oxidation gene program transcriptomic expression and expression of all TFs (Govaere et al.). (N) Maximally prioritized TFs inferred as driving (top 5) or repressing (bottom 5) beta-oxidation gene program. (O) Summary of inferred regulatory relationships between TFs chosen for experimental validation and each stress adaptation-associated gene program.

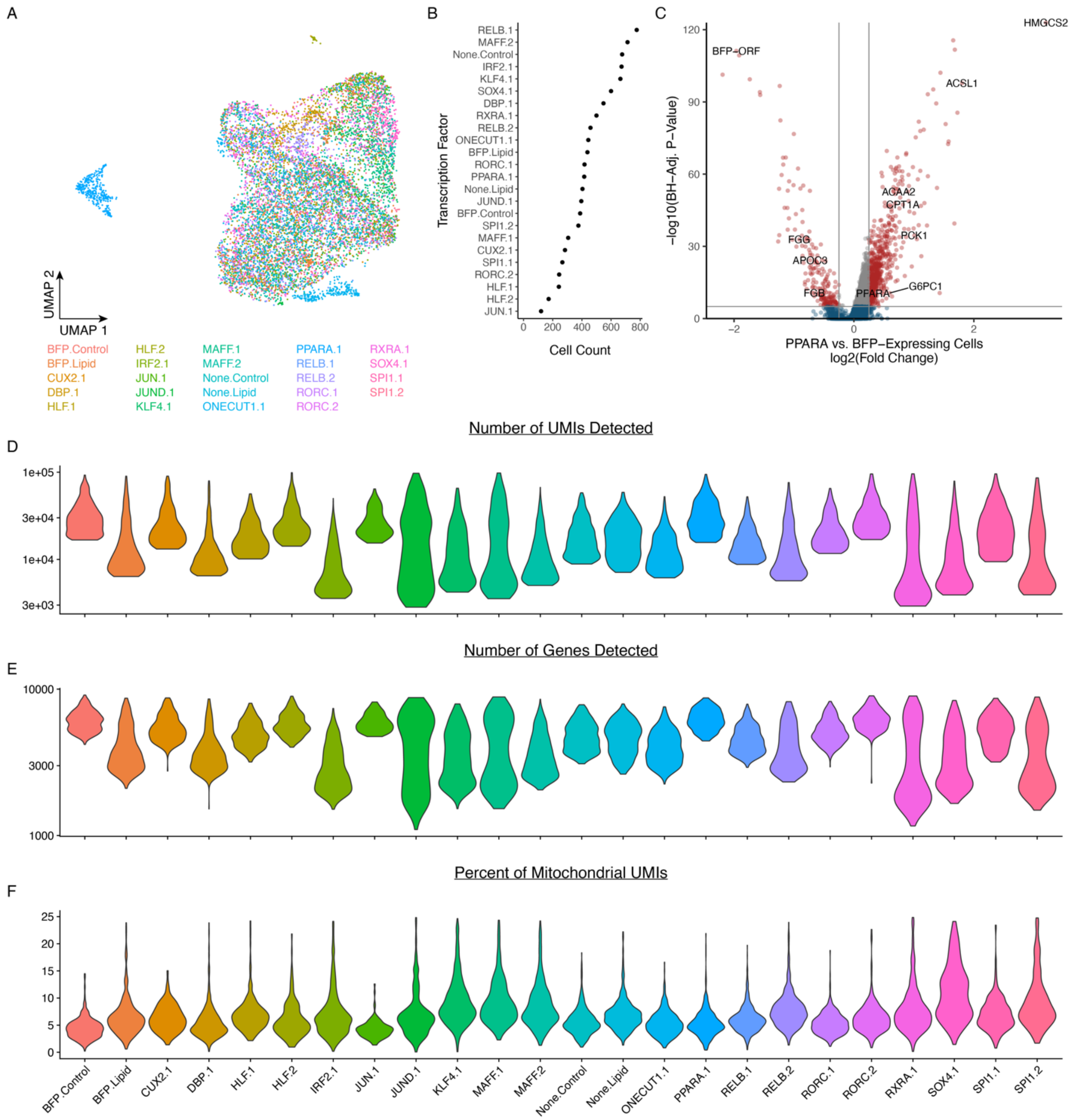


Supplementary Figure 12: Data overview and QC metrics of arrayed human genetic perturbation (lentiviral TF overexpression) scRNA-seq. (A) UMAP visualization of arrayed lentiviral TF overexpression scRNA-seq data. (B) Number of cells passing QC per overexpressed TF. (C) Differentially-expressed genes between PPARA-overexpressing vs. BFP-overexpressing cells in lipid-rich media. (D) Number of detected UMIs, split by TF. (E) Number of detected genes, split by TF. (F) Percent of mitochondrial UMIs, split by TF.

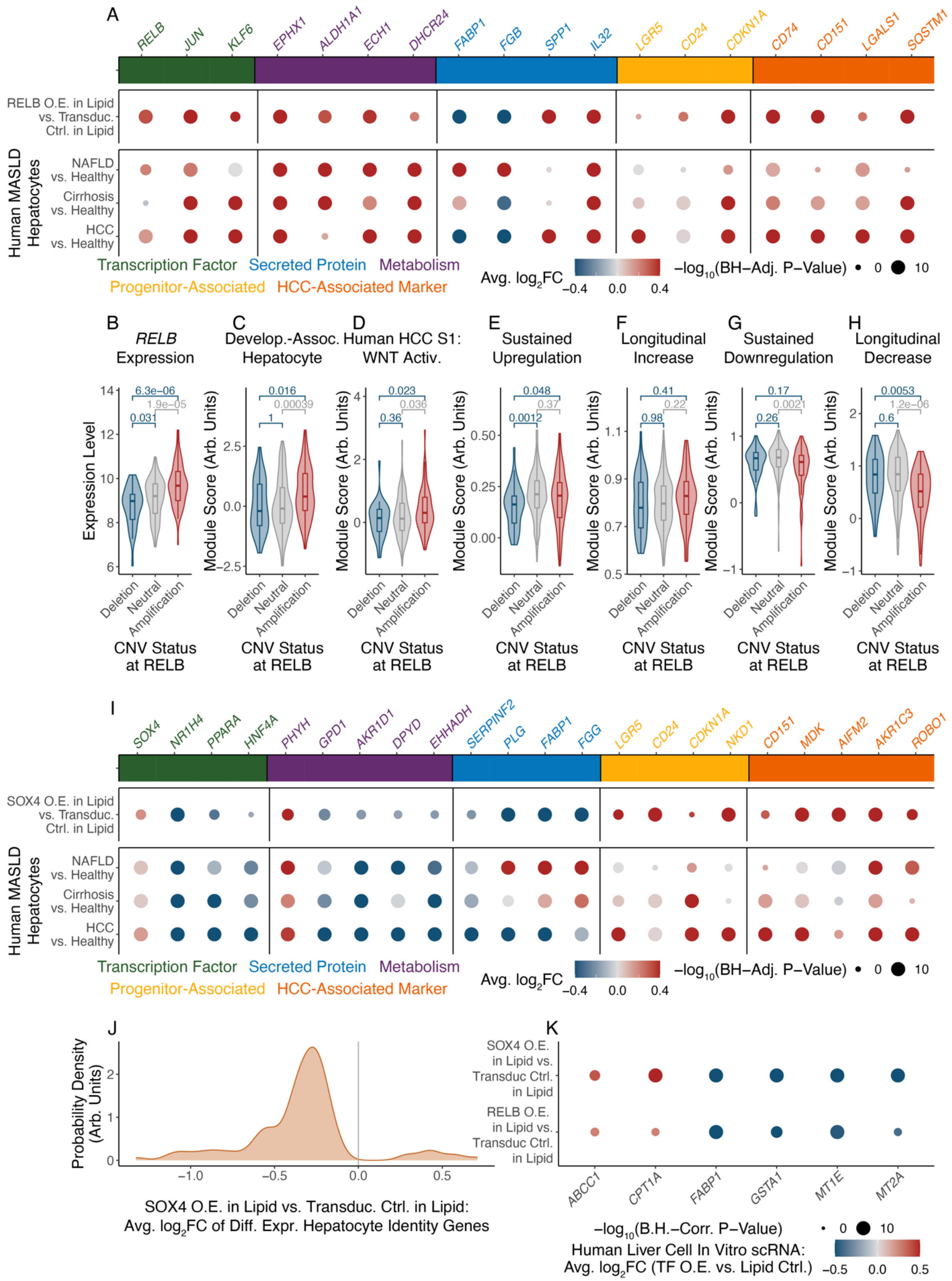
A
RELB
JUN
KLF6
EPHX1
ALDH1A1
ECH1
DHCR24
FABP1
FGB
SPP1
IL32
LGR5
CD24
CDKN1A
CD74
CD151
LGALS1
SQSTM1
RELB O.E. in Lipid vs. Transduc. Ctrl. in Lipid
Human MASLD Hepatocytes
NAFLD vs. Healthy
Cirrhosis vs. Healthy
HCC vs. Healthy
Transcription Factor
Secreted Protein
Metabolism
Progenitor-Associated
HCC-Associated Marker
Avg. log2FC
−0.4
0.0
0.4
−log10(BH−Adj. P−Value)
0
10
B
RELB Expression
C
Develop.-Assoc. Hepatocyte
D
Human HCC S1: WNT Activ.
E
Sustained Upregulation
F
Longitudinal Increase
G
Sustained Downregulation
H
Longitudinal Decrease
Expression Level
Module Score (Arb. Units)
6.3e−06
1.9e−05
0.031
0.016
0.00039
1
0.023
0.036
0.36
0.048
0.37
0.0012
0.41
0.22
0.98
0.17
0.0021
0.26
0.0053
1.2e−06
0.6
Deletion
Neutral
Amplification
CNV Status at RELB
I
SOX4
NR1H4
PPARA
HNF4A
PHYH
GPD1
AKR1D1
DPYD
EHHADH
SERPINF2
PLG
FABP1
FGG
LGR5
CD24
CDKN1A
NKD1
CD151
MDK
AIFM2
AKR1C3
ROBO1
SOX4 O.E. in Lipid vs. Transduc. Ctrl. in Lipid
Human MASLD Hepatocytes
NAFLD vs. Healthy
Cirrhosis vs. Healthy
HCC vs. Healthy
Transcription Factor
Secreted Protein
Metabolism
Progenitor-Associated
HCC-Associated Marker
Avg. log2FC
−0.4
0.0
0.4
−log10(BH−Adj. P−Value)
0
10
J
Probability Density (Arb. Units)
−1.0
−0.5
0.0
0.5
SOX4 O.E. in Lipid vs. Transduc. Ctrl. in Lipid: Avg. log2FC of Diff. Expr. Hepatocyte Identity Genes
K
SOX4 O.E. in Lipid vs. Transduc Ctrl. in Lipid
RELB O.E. in Lipid vs. Transduc Ctrl. in Lipid
ABCC1
CPT1A
FABP1
GSTA1
MT1E
MT2A
−log10(B.H.−Corr. P−Value)
0
10
Human Liver Cell In Vitro scRNA: Avg. log2FC (TF O.E. vs. Lipid Ctrl.)
−0.5
0.0
0.5

**Supplementary Figure 13: Contextualization of regulatory effects of RELB and SOX4 on hepatocytes' metabolic adaptation programs.** (A) Gene expression change induced by RELB overexpression in lipid-rich media (compared to BFP transduction control in lipid-rich media) (top row), or gene expression change with MASLD/cirrhosis/HCC progression in human patients' hepatocytes (Filliol et al. human snRNA-seq dataset). (B-H) TCGA HCC samples stratified by CNV status at RELB gene locus, with expression of (B) RELB itself, (C) Development-Associated Hepatocyte gene program, (D) Human HCC S1: WNT Activation gene program, (E) Sustained Upregulation gene program, (F) Longitudinal Increase gene program, (G) Sustained Downregulation gene program, (H) Longitudinal Decrease gene program. (I) Gene expression change induced by SOX4 overexpression in lipid-rich media (compared to BFP transduction control in lipid-rich media) (top row), or gene expression change with MASLD/cirrhosis/HCC progression in human patients' hepatocytes (Filliol et al. human snRNA-seq dataset). (J) Distribution of expression fold-changes among genes undergoing significant changes with SOX4 overexpression in lipid-rich media (compared to BFP transduction control in lipid-rich media). (K) Fold-change of genes involved in lipid handling and oxidative stress response with overexpression of SOX4 or RELB in lipid-rich media (compared to BFP transduction control in lipid-rich media). All p-values calculated using Mann-Whitney U test.

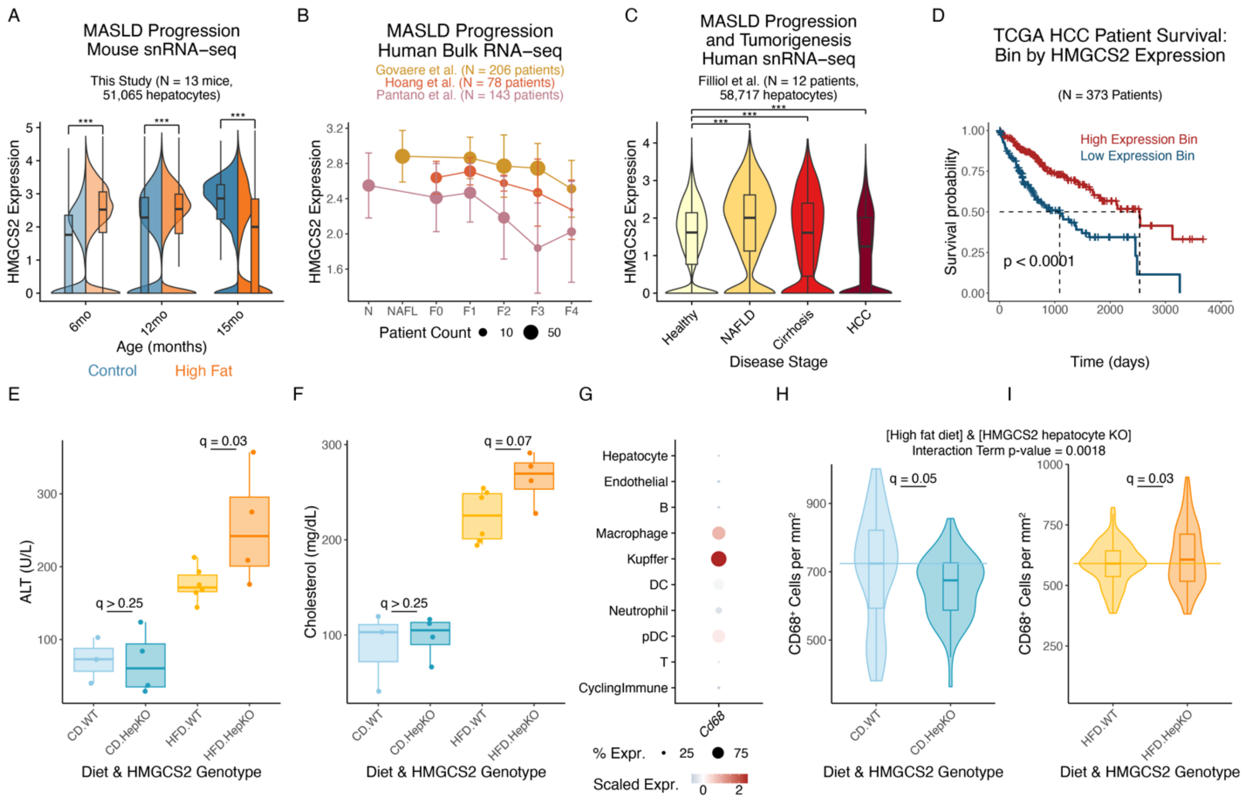


Supplementary Figure 14: Characterization of HMGCS2 with chronic metabolic stress adaptation and hepatocyte-specific knockout cohort. (A) *Hmgcs2* expression during chronic metabolic stress natural progression in mice (snRNA-seq). (B) *HMGCS2* expression during MASLD in human patient cohorts (bulk RNA-seq). (C) *HMGCS2* expression during MASLD/cirrhosis/HCC in human patient cohorts (snRNA-seq). (D) Stratification of TCGA HCC patient survival, based on expression of *HMGCS2*. (E) ALT levels in HMGCS2 knockout cohort. (F) Cholesterol levels in HMGCS2 knockout cohort. (G) Expression of *Cd68* across cell types in mice (scRNA-seq). (H-I) IHC staining for *in situ* abundance of CD68+ cells in tissue sections from control diet (H) or high fat diet (I) validation cohort mice. Survival outcome p-values calculated with log-rank test. ALT, cholesterol, and CD68$^+$ cell abundance statistics calculated using Student's t-test with Benjamini-Hochberg multiple-testing correction. All other p-values calculated using Mann-Whitney U test.

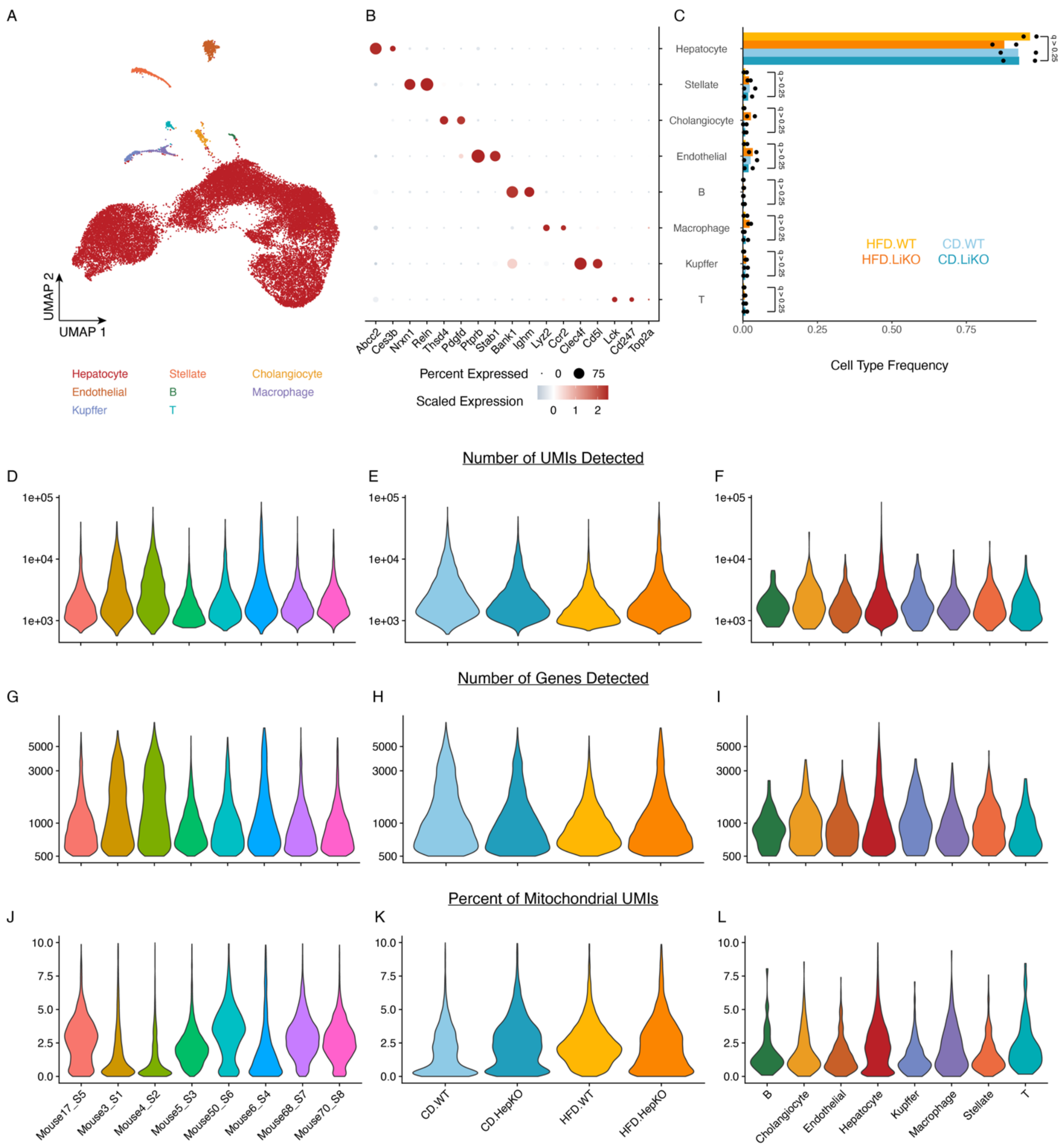


<u>Supplementary Figure 15: Data overview and QC metrics of frozen-tissue snRNA-seq on hepatocyte-specific HMGCS2 knockout cohort.</u> (A) UMAP visualization of frozen-tissue tumor snRNA-seq data in hepatocyte-specific HMGCS2 knockout cohort. (B) Marker gene dotplot visualization and (C) compositional abundance of cell types. (D-F) Number of detected UMIs split by (D) mouse, (E) diet and Hmgcs2 genotype status, or (F) cell type. (G-I) Number of detected genes split by (G) mouse, (H) diet and HMGCS2 genotype status, or (I) cell type. (J-L) Percent of

mitochondrial UMIs split by (J) mouse, (K) diet and tumor status, or (L) cell type. Survival outcome p-values calculated with log-rank test; all other p-values calculated using Mann-Whitney U test.
